# Dynamic and selective engrams emerge with memory consolidation

**DOI:** 10.1101/2022.03.13.484167

**Authors:** Douglas Feitosa Tomé, Ying Zhang, Tomomi Aida, Sadra Sadeh, Dheeraj S. Roy, Claudia Clopath

## Abstract

Episodic memories are encoded by sparse populations of neurons activated during an experience.^1^ These neural ensembles constitute memory engrams that are both necessary and sufficient for inducing recall even long after memory acquisition.^2^ This suggests that following encoding, engrams are stabilized to reliably support memory retrieval. However, little is known about the temporal evolution of engrams over the course of memory consolidation or how it impacts mnemonic properties. Here we employed computational and experimental approaches to examine how the composition and selectivity of engrams change with memory consolidation. We modeled engram cells using a spiking recurrent neural network that yielded three testable predictions: memories transition from unselective to selective as neurons are removed from and added to the engram, inhibitory activity during recall is essential for memory selectivity, and inhibitory synaptic plasticity during memory consolidation is critical for engrams to become selective. Using the Cal-Light system to tag activated neurons *in vivo* with high spatiotemporal precision^3^ as well as optogenetic and chemogenetic techniques, we conducted contextual fear conditioning experiments that supported each of our model’s predictions. Our results reveal that engrams are dynamic even within hours of memory consolidation and that changes in engram composition mediated by inhibitory synaptic plasticity are crucial for the emergence of memory selectivity. These findings challenge classical theories of stable memory traces and point to a close link between engram state and memory expression.

## Main

A growing body of evidence has shown that neurons activated by an experience have a prominent role in memory.^2^ Specifically, loss-of-function and gain-of-function studies have demonstrated that ablating neurons activated during learning disrupts memory retrieval^4^ whereas artificially reactivating these neurons elicits memory recall even in the absence of retrieval cues.^5^ Therefore, learning-activated neurons are a cellular substrate for memory storage and retrieval and they constitute engram cells. The stability of engrams following memory encoding, however, remains an open question. In particular, there are two competing hypotheses regarding the effect of memory consolidation on the post-learning evolution of engrams. First, engrams may be stabilized as a result of memory consolidation (i.e., stable engrams) in line with the crucial role of encoding-activated engram cells in subsequent memory retrieval.^2^ Second, the relatively low overlap between ensembles of neurons activated during both learning and recall (∼ 10 to 40%) raises the possibility that engrams may change over the course of memory consolidation with neurons “dropping out of” or “dropping into” the engram (i.e., dynamic engrams).^2^ Critically, knowledge of the temporal profile of engrams may elucidate how engram composition is related to mnemonic properties such as memory selectivity — a feature essential for adaptive behavior.^6^

### Memory consolidation reshapes engrams

We used a computational model to probe the evolution of engram cells and their selectivity with memory consolidation. Specifically, our spiking neural network model consisted of a stimulus population that projected to the hippocampus (Fig. 1a). Feedforward and recurrent excitatory synapses onto excitatory neurons exhibited short- and long-term plasticity while inhibitory synapses onto excitatory neurons displayed inhibitory plasticity. Long-term excitatory synaptic plasticity combined a Hebbian term consisting of triplet spike-timing-dependent plasticity (STDP)^7^ and non-Hebbian terms including heterosynaptic plasticity^8^ and transmitter-induced plasticity^9^ as proposed previously.^10^ Inhibitory synaptic plasticity took the form of a network activity-based STDP mechanism^10^ whose primary goal was to control network activity levels^11^ (see Methods for a detailed description of the model). Our network was initially trained by presenting a training stimulus to simulate an episodic memory task (Fig. 1b-c). We then identified training-activated engram cells by examining which neurons were selectively activated in response to the training stimulus (i.e., average stimulus-evoked firing rate above the threshold *ζ^thr^* = 10 Hz in the last 60 s of the training phase). Subsequently, the network underwent a consolidation period when the training stimulus was reactivated in line with previous experimental reports that learning-activated sensory neurons are reactivated during post-encoding sleep.^12^ At regular intervals throughout the consolidation phase (i.e., consolidation time = 0, 1, …, 24 h), we investigated the state of the engram by presenting the training stimulus in a probing phase and identifying engram cells at that time point in a manner analogous to the one in the training phase. Probing-activated engram cells then represented the current state of the engram following memory consolidation. Lastly, we presented partial cues of either the training stimulus or a novel, unseen stimulus in the recall phase (Fig. 1b-c). In particular, we conducted a total of four separate recall sessions (i.e., one for the training stimulus and one for each of the three novel stimuli) after every sampled consolidation interval. This allowed us to evaluate the ability of the network to selectively recall the encoded memory only when cues of the training stimulus were presented. Note that the overlap between the training stimulus (i.e., square) and each novel stimulus (i.e., circle, pentagon, and hexagon) varies with the circle having the highest overlap (Extended Data Fig. 1a).

**Fig. 1.**
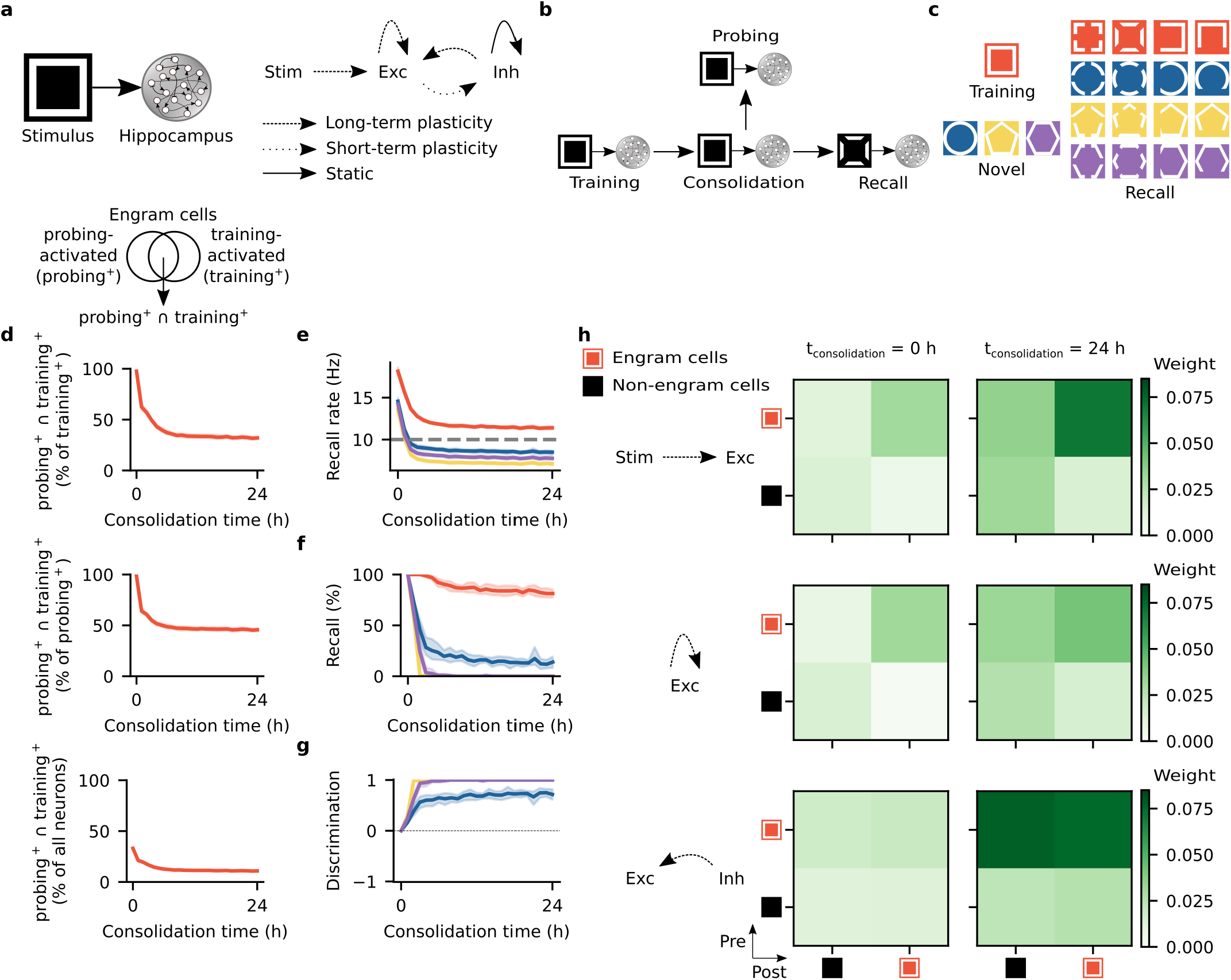
Memory consolidation renders engrams dynamic and selective. **a**, Schematic of computational model. Left: Stimulus population (Stim) and hippocampus network with excitatory (Exc) and inhibitory (Inh) neurons. Right: plasticity of feedforward and recurrent synapses (see Methods). **b**, Schematic of simulation protocol. **c**, Schematic of training and novel stimuli with corresponding partial cues for recall. **d-g**, Means and 99% confidence intervals shown. *n* = 10. **d**, Post-encoding evolution of engram cells. Ensemble overlap between engram cells activated during both probing and training as a fraction of training-activated engram cells (top), probing-activated engram cells (middle), and all neurons in the network (bottom). **e**, Firing rate of engram cells averaged across all cue presentations during recall as a function of consolidation time. Memory recall was tested with cues of the training and novel stimuli (see Methods). Dashed line indicates threshold *ζ^thr^* = 10 Hz for engram cell activation. Color denotes stimulus as in **c**. **f**, Memory recall as a function of consolidation time. Color denotes stimulus as in **c**. **g**, Discrimination index between recall evoked by cues of the training stimulus and individual novel stimuli as a function of consolidation time (see Methods). Color denotes stimulus as in **c**. **h**, Mean weight strength of plastic synapses clustered according to engram cell status (i.e., engram and non-engram cells). Top: feedforward excitatory synapses onto excitatory neurons. Middle: recurrent excitatory synapses onto excitatory neurons. Bottom: recurrent inhibitory synapses onto excitatory neurons. Left: at the end of the training phase. Right: after 24 hours of consolidation. Representative trial shown.

Our simulation results showed that memory consolidation reorganized engrams in our model with neurons being removed from and added to the engram (Fig. 1d). We tracked the fraction of training-activated engram cells that remained part of the engram over the course of memory consolidation by computing the ensemble overlap between probing-activated engram cells and training-activated engram cells as a fraction of training-activated engram cells. Given that this ratio decreased substantially as consolidation progressed, our model predicted that a large fraction of training-activated engram cells are removed from the engram and no longer actively encode the memory acquired during training (i.e., they stop being activated by the training stimulus). To determine what fraction of probing-activated engram cells were originally training-activated engram cells, we computed the ensemble overlap between probing-activated engram cells and training-activated engram cells as a fraction of probing-activated engram cells. This quantity also significantly decreased with memory consolidation. Thus, our model predicted that neurons that were not activated during training are recruited into the engram and start actively encoding the underlying memory (i.e., they become responsive to the training stimulus). Additionally, we found that the ensemble overlap between probing-activated engram cells and training-activated engram cells as a fraction of all neurons in the network slowly decreased over the course of consolidation. This indicated that the ensemble of probing-activated engram cells gradually shrank since the ensemble of training-activated engram cells was fixed in any given trial (Extended Data Fig. 1b). Furthermore, our network exhibited a high overlap between probing-activated engram cell ensembles identified in two consecutive hours but only a very small fraction of training-activated engram cells remained part of the engram in every sampled consolidation interval (Extended Data Fig. 1c). Therefore, our model predicted that memory engrams are highly dynamic with neurons being removed from and added to the engram over the course of memory consolidation. Dynamic inhibitory engrams also emerged in our network model but with a more moderate turnover rate (Extended Data Fig. 2a-c).

In addition, engrams in our model were initially unselective but became selective over the course of memory consolidation (Fig. 1e-g). To examine engram selectivity, we first defined recall rate as the cue-evoked population firing rate of engram cells averaged across all cue presentations in the recall phase (see Methods). We saw that the recall rate elicited by cues of the training and novel stimuli were above the threshold *ζ^thr^* = 10 Hz for engram cell activation immediately following training but as memory consolidation progressed only the recall rate of cues of the training stimulus remained above the activation threshold (Fig. 1e). We also specifically measured memory recall by computing the fraction of cue presentations in the recall phase that activated the engram cell ensemble (an engram cell ensemble was considered activated when its average firing rate was above the activation threshold during the presentation of a cue, see Methods). This recall metric revealed equal memory recall levels for the training and novel stimuli immediately following training but with memory consolidation only the recall of the training stimulus remained at a high level (Fig. 1f, Extended Data Fig. 3). We then defined a discrimination index as the difference between recall of the training stimulus and recall of a novel stimulus divided by their sum to provide a normalized measure of memory selectivity (see Methods). This discrimination index showed that our network developed the ability to distinguish between the training and novel stimuli as a result of memory consolidation (Fig. 1g). The slightly lower post-consolidation discrimination index of one of the novel stimuli (i.e., circle) can be attributed to its higher overlap with the training stimulus as mentioned previously (Extended Data Fig. 1a). Thus, our model predicted that engrams are initially encoded in an unselective state but later become selective as its composition changes with memory consolidation. This is consistent with a recent study showing that memory selectivity in conditioned taste aversion emerges in the timescale of hours.^13^ Note also that even though the fraction of probing-activated engram cells reactivated during recall was higher for the training stimulus relative to the novel stimuli, this ratio was only ∼ 50% for the training stimulus after memory consolidation (Extended Data Fig. 1d). This suggested that our network model performed an operation akin to pattern separation — a process that has been ascribed to the dentate gyrus (DG) and CA3 of the hippocampus.^14^ Lastly, inhibitory engrams remained unselective in our simulations (Extended Data Fig. 2d-g).

The changes in engram composition and selectivity observed in our model were associated with ongoing synaptic plasticity during memory consolidation (Fig. 1h). Feedforward synapses from training stimulus neurons (i.e., sensory engram cells, see Methods) onto hippocampal engram cells were strengthened over the course of memory consolidation and, consequently, the synaptic coupling between the stimulus population and the hippocampus network was increased. Recurrent excitatory synapses among engram cells also experienced a modest gain in synaptic efficacy. Notably, inhibitory synapses from inhibitory engram cells onto both engram and non-engram cells were strongly potentiated throughout memory consolidation. This indicated that a number of training-activated engram cells were forced out of the engram due to strong inhibition and only neurons highly responsive to the training stimulus remained in the engram in line with our previous analysis (Fig. 1d). Inhibitory neurons also controlled the overall activity of excitatory neurons in the network through inhibitory synaptic plasticity (Extended Data Fig. 2h).

To investigate the contribution of synaptic plasticity to the engram dynamics in our model, we performed several loss-of-function and gain-of-function manipulations. First, we blocked the reactivation of the training stimulus during memory consolidation and found that this altered the temporal profile of engrams and prevented them from becoming selective (Extended Data Fig. 4). These effects were associated with reduced potentiation of inhibitory synapses onto engram cells (compare Extended Data Fig. 4i to Fig. 1h). Previous experiments demonstrated that sleep-specific inhibition of learning-activated sensory neurons disrupts memory selectivity^12^ and, hence, our model was consistent with these findings and it predicted underlying mechanisms. Second, blocking long-term potentiation (LTP) during memory consolidation almost completely eliminated engram cell turnover after a steady state was reached and it also impaired memory recall relative to the control case (Extended Data Fig. 5). Reduced feedforward and recurrent excitatory synaptic weights due to LTP blockage led to engram stabilization and impaired recall (compare Extended Data Fig. 5h to Fig. 1h). These results were in line with a recent study showing that memory recall is impaired when LTP is optically erased selectively during sleep.^15^ Third, we separately blocked the Hebbian and non-Hebbian forms of long-term excitatory synaptic plasticity in our model and verified that each was essential for memory encoding and consolidation (Extended Data Fig. 6). Specifically, blocking triplet STDP disrupted the potentiation of recurrent synapses among engram cells leading to impaired memory encoding. On the other hand, blocking heterosynaptic plasticity enabled excessive potentiation of excitatory synapses to the point where even robust recruitment of inhibition could not support memory selectivity. Conversely, blocking transmitter-induced plasticity impaired memory selectivity due to reduced potentiation of inhibitory synapses. The previous results were consistent with a mean-field analysis showing that this combination of plasticity mechanisms can support stable memory formation and recall.^10^ Furthermore, blocking training-activated engram cells after a consolidation period of 24 h prevented memory recall (Extended Data Fig. 1e) whereas artificially reactivating them in the absence of retrieval cues was able to elicit recall (Extended Data Fig. 1f) in a manner consistent with previous experimental findings^4, 5^ and despite the dynamic nature of engrams in our simulations. Thus, our model was able to reconcile the prominent role of training-activated engram cells in memory storage and retrieval with dynamic memory engrams. Finally, we found that an alternative inhibitory synaptic plasticity formulation yielded engram dynamics analogous to those in our original network (compare Extended Data Fig. 7a-h to Fig. 1d-h and Extended Data Fig. 1b-d). This suggested that the dynamic and selective engrams predicted by our model are not a product of a specific form of inhibitory plasticity but a consequence of memory encoding and consolidation in inhibition-stabilized plastic networks in general.

### Dynamic and selective fear memory engrams

To test our model’s prediction that memories switch from unselective to selective as the composition of the underlying engram changes, we used a contextual fear conditioning paradigm. Specifically, mice were fear conditioned in a training context and following a delay period memory recall was assessed at first in a neutral context and subsequently in the training context (Fig. 2a, see Methods). Freezing levels in the training and neutral contexts were comparable shortly after fear training (i.e., delay time = 1 and 5 h) but they differed for longer delay periods (i.e., delay time = 12, 18 and 24 h) (Fig. 2b). By defining a memory discrimination index for the freezing behavior in the training and neutral contexts analogous to the one for recall in our simulations (see Methods), we saw that fear memory selectivity emerged after a delay of 12 h following training (Fig. 2c). These findings were consistent with our modeling predictions (compare Fig. 2b-c to Fig. 1f-g) as well as with recent reports that conditioned taste aversion memories become selective in the timescale of hours.^13^ To track the evolution of fear memory engrams, we used the Cal-Light system to tag hippocampal DG neurons active during fear training with EGFP^3^ and we labeled neurons active during recall tests using *c-Fos* staining (Fig. 2a and Extended Data Fig. 8a left panel, see Methods). We then observed that when mice were returned to the training context for recall each of the following ratios decreased over time: ensemble overlap between neurons activated during both recall and training (*c-Fos*^+^ ∩ EGFP^+^) as a fraction of training-activated neurons (EGFP^+^), recall-activated neurons (*c-Fos*^+^), and neuronal cell count (DAPI^+^) (Fig. 2d). The decline in the level of recall-induced reactivation of training-activated neurons was consistent with our prediction that only a fraction of training-activated engram cells remain in the engram as memory consolidation progresses (Fig. 1d). The drop in the proportion of recall-activated neurons that were also active during training was in line with our prediction that new neurons are recruited into the engram after memory encoding (Fig. 1d). The decrease in the ensemble overlap between recall- and training-activated neurons relative to neuronal cell count was also consistent with our modeling results (Fig. 1d). *c-Fos*^+^ ∩ EGFP^+^ as a fraction of EGFP^+^, *c-Fos*^+^ and DAPI^+^ also dropped over time when mice were placed in the neutral context after fear training (Fig. 2d). To determine baseline levels for these ensemble overlap ratios, we measured them when mice were placed in their home cage following conditioning (Fig. 2d). Critically, ensemble overlap levels for recall tests in the training and neutral contexts were both above the home cage baseline at delay time = 5 h but only recall in the training context elicited ensemble overlap ratios above baseline at delay time = 12 h (Fig. 2d). This coincided with the emergence of fear memory selectivity (Fig. 2c) and was consistent with our network simulations (Fig. 1e/g). Notably, *c-Fos*^+^ ∩ EGFP^+^ as a fraction of EGFP^+^ and DAPI^+^ for recall in the training context remained above the home cage baseline at delay time = 24 h, but not *c-Fos*^+^ ∩ EGFP^+^ as a fraction of *c-Fos*^+^. This could indicate that the rate of engram cell turnover at this point was elevated such that it would require highly precise measurements to capture differences in the fraction of recall-activated neurons that were also active during encoding when mice are placed in the training context versus the home cage. Altogether, our experimental results supported our model’s prediction that memory consolidation leads to dynamic and selective engrams.

**Fig. 2.**
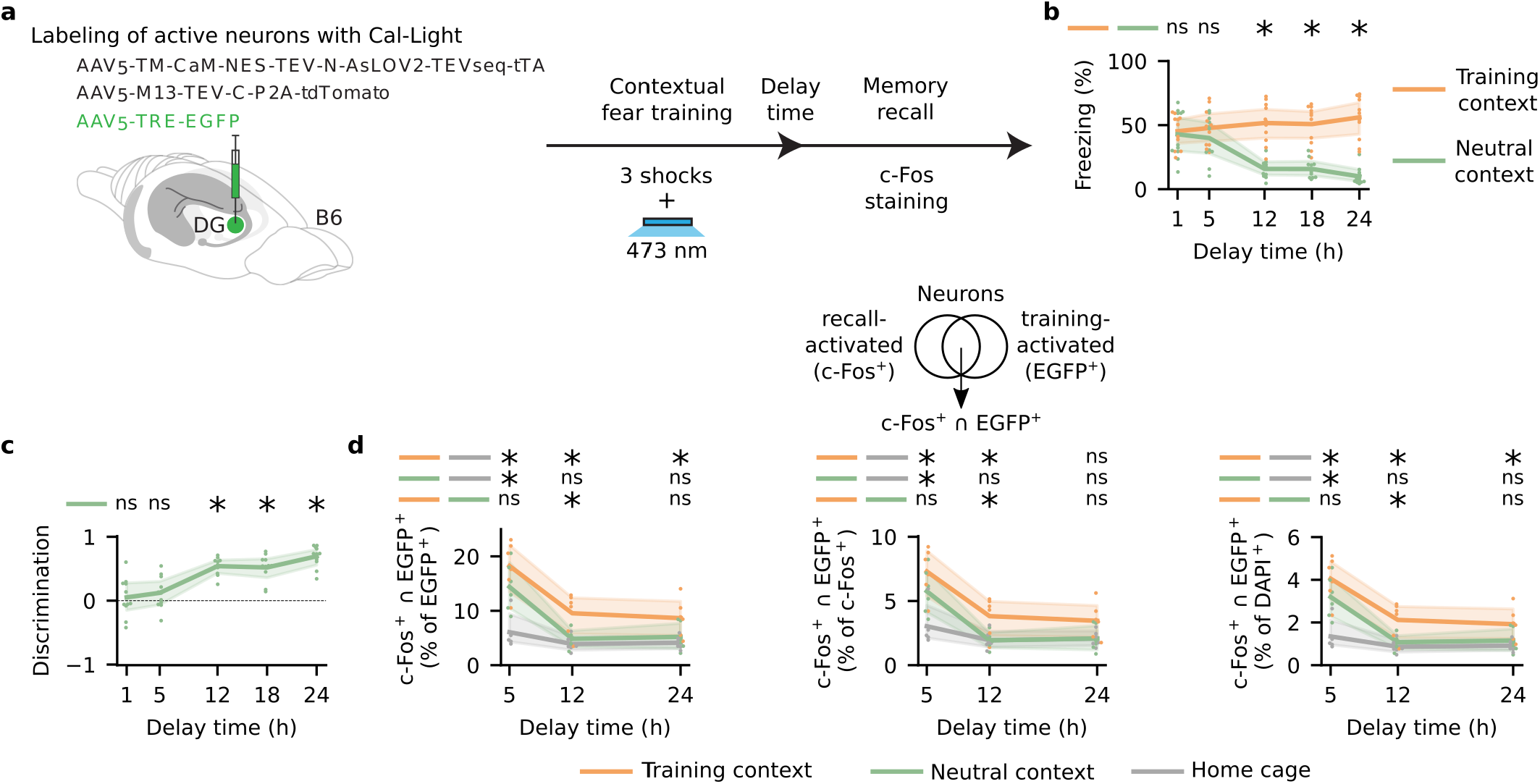
Dynamic and selective engrams encode contextual fear memory. **a**, Schematic of experimental protocol. **b-d**, Means and 99% confidence intervals shown. **b**, Freezing levels during memory recall as a function of delay time. Memory recall was tested in the training and neutral contexts (see Methods). Two-sided Wilcoxon signed-rank test between freezing in the two contexts. delay = 1 h: *W* = 29.0, *P* = 0.764648. delay = 5 h: *W* = 19.0, *P* = 0.240234. delay = 12 h: *W* = 0.0, *P* = 0.000977. delay = 18 h: *W* = 0.0, *P* = 0.000977. delay = 24 h: *W* = 0.0, *P* = 0.000977. *n* = 11 mice per group. **c**, Discrimination index between freezing levels in the training and neutral contexts as a function of delay time (see Methods). Two-sided one-sample Wilcoxon signed-rank test against discrimination = 0. delay = 1 h: *W* = 29.0, *P* = 0.764648. delay = 5 h: *W* = 17.0, *P* = 0.174805. delay = 12 h: *W* = 0.0, *P* = 0.000977. delay = 18 h: *W* = 0.0, *P* = 0.000977. delay = 24 h: *W* = 0.0, *P* = 0.000977. *n* = 11 mice per group. **d**, Ensemble overlap between neurons activated during both recall and training (*c-Fos*^+^ ∩ EGFP^+^) as a fraction of training-activated neurons (EGFP^+^) (left), recall-activated neurons (*c-Fos*^+^) (middle), and neuronal cell count (DAPI^+^) (right). Memory recall was tested in the training context, neutral context, and home cage (see Methods). One-way ANOVA followed by Tukey’s HSD post hoc test. *c-Fos*^+^ ∩ EGFP^+^ (% of EGFP^+^): for delay = 5 h, *F*_2,15_ = 13.883272, *P* = 0.000387 (ANOVA), *P* = 0.001000 (training context vs. home cage), *P* = 0.008067 (neutral context vs. home cage), and *P* = 0.264460 (training context vs. neutral context); for delay = 12 h, *F*_2,15_ = 8.691289, *P* = 0.003114 (ANOVA), *P* = 0.003855 (training context vs. home cage), *P* = 0.778343 (neutral context vs. home cage), and *P* = 0.014340 (training context vs. neutral context); and for delay = 24 h, *F*_2,15_ = 5.000676, *P* = 0.021675 (ANOVA), *P* = 0.021735 (training context vs. home cage), *P* = 0.741921 (neutral context vs. home cage), and *P* = 0.085645 (training context vs. neutral context). *c-Fos*^+^ ∩ EGFP^+^ (% of *c-Fos*^+^): for delay = 5 h, *F*_2,15_ = 9.495859, *P* = 0.002165 (ANOVA), *P* = 0.001703 (training context vs. home cage), *P* = 0.037383 (neutral context vs. home cage), and *P* = 0.295178 (training context vs. neutral context); for delay = 12 h, *F*_2,15_ = 6.582162, *P* = 0.00887 (ANOVA), *P* = 0.017565 (training context vs. home cage), *P* = 0.900000 (neutral context vs. home cage), and *P* = 0.017088 (training context vs. neutral context); and for delay = 24 h, *F*_2,15_ = 3.427944, *P* = 0.059418 (ANOVA), *P* = 0.089977 (training context vs. home cage), *P* = 0.900000 (neutral context vs. home cage), and *P* = 0.093594 (training context vs. neutral context). *c-Fos*^+^ ∩ EGFP^+^ (% of DAPI^+^): for delay = 5 h, *F*_2,15_ = 13.888714, *P* = 0.000386 (ANOVA), *P* = 0.001000 (training context vs. home cage), *P* = 0.008062 (neutral context vs. home cage), and *P* = 0.264231 (training context vs. neutral context); for delay = 12 h, *F*_2,15_ = 8.682464, *P* = 0.003127 (ANOVA), *P* = 0.003870 (training context vs. home cage), *P* = 0.778756 (neutral context vs. home cage), and *P* = 0.014377 (training context vs. neutral context); and for delay = 24 h, *F*_2,15_ = 5.00044, *P* = 0.021678 (ANOVA), *P* = 0.021749 (training context vs. home cage), *P* = 0.742354 (neutral context vs. home cage), and *P* = 0.085571 (training context vs. neutral context). *n* = 6 mice per group. **b-d**, *: *P* < 0.05; ns: not significant.

### Inhibition is crucial for memory selectivity

We next examined the role of inhibition in memory selectivity using our network model. We blocked inhibitory neurons during memory recall and this disrupted selectivity in our simulations (Fig. 3a-d, see with alternative inhibitory synaptic plasticity Extended Data Fig. 7i). Previous experiments showed that inhibiting DG cholecystokinin-expressing (CCK^+^) interneurons during recall 24 h after contextual fear conditioning impairs memory selectivity but inhibiting DG parvalbumin-expressing (PV^+^) interneurons during recall at the same time point has no effect on selectivity.^6^ This was associated to an enhanced post-training inhibitory synaptic input from DG CCK^+^ but not PV^+^ interneurons onto training-activated engram cells.^6^ To capture this difference between interneuron types in mediating memory selectivity, we expanded our network model to include both CCK^+^ and PV^+^ interneurons (Extended Data Fig. 9a). CCK^+^ but not PV^+^ synapses onto excitatory neurons were plastic consistent with experimental reports.^6^ Dynamic and selective engrams emerged in the expanded model in a manner analogous to our original network (compare Extended Data Fig. 9b-i to Fig. 1d-h and Extended Data Fig. 1b-d). In addition, blocking CCK^+^ interneurons during recall disrupted memory selectivity but blocking PV^+^ interneurons during recall had no such impact (Extended Data Fig. 9j-k) in line with experimental results.^6^ Note that blocking either CCK^+^ or PV^+^ interneurons prior to the emergence of selectivity (e.g., immediately following training) had a negligible effect on memory recall and discrimination (compare Extended Data Fig. 9j-k to f-g). Thus, our model predicted that inhibitory neurons — and specifically DG CCK^+^ interneurons — are essential for expressing memory selectivity once engrams have become selective. To test this prediction, we used CCK-Cre mice to optogenetically inhibit DG CCK^+^ interneurons during fear memory recall (Fig. 3e and Extended Data Fig. 8a middle panel, see Methods). Mice in the CCK^+^ inhibition group were unable to discriminate between the training and neutral contexts at all tested delay times while those in the control group displayed memory selectivity with delay times of 12 and 24 h (Fig. 3f and Extended Data Fig. 8b, compare to Fig 2b-c). Therefore, our experimental findings demonstrated that DG CCK^+^ interneurons are required for memory discrimination and, hence, they supported our model’s prediction that inhibitory activity during recall is critical for engram selectivity.

**Fig. 3.**
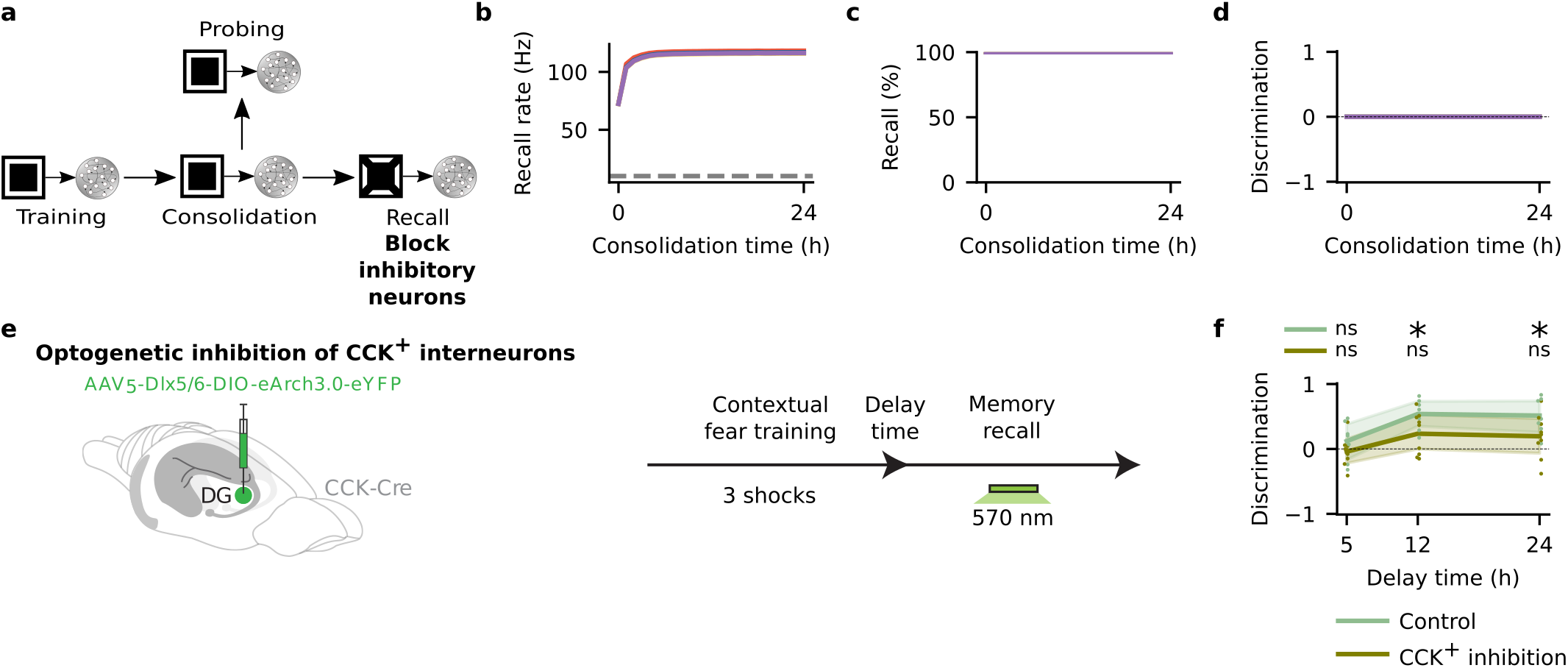
Inhibitory activity during recall is critical for memory selectivity. **a**, Schematic of simulation protocol with blockage of inhibitory neurons during recall. **b-d**, Means and 99% confidence intervals shown. *n* = 10. **b**, Firing rate of engram cells averaged across all cue presentations during recall in **a** as a function of consolidation time (see Methods). Dashed line indicates threshold *ζ^thr^* = 10 Hz for engram cell activation. Color denotes stimulus as in Fig. 1c. **c**, Memory recall in **a** as a function of consolidation time. Color denotes stimulus as in Fig. 1c. **d**, Discrimination index between recall evoked by cues of the training stimulus and individual novel stimuli in **a** as a function of consolidation time (see Methods). Color denotes stimulus as in Fig. 1c. **e**, Schematic of experimental protocol with optogenetic inhibition of DG CCK^+^ interneurons. **f**, Discrimination index between freezing levels in the training and neutral contexts in **e** as a function of delay time (see Methods). Means and 99% confidence intervals shown. Two-sided one-sample Wilcoxon signed-rank test against discrimination = 0. Control (*n* = 7 mice per group): for delay = 5 h, *W* = 9.0, *P* = 0.46875; for delay = 12 h, *W* = 0.0, *P* = 0.015625; and for delay = 24 h, *W* = 0.0, *P* = 0.015625. CCK^+^ inhibition (*n* = 9 mice per group): for delay = 5 h, *W* = 17.0, *P* = 0.570313; for delay = 12 h, *W* = 9.0, *P* = 0.128906; and for delay = 24 h, *W* = 9.0, *P* = 0.128906. *: *P* < 0.05; ns: not significant.

### Inhibitory plasticity molds selective engrams

Given the essential role of inhibition in the expression of memory selectivity, we investigated the contribution of post-encoding inhibitory synaptic plasticity to the emergence of selective engrams. To that end, we blocked inhibitory synaptic plasticity exclusively during memory consolidation in our network simulations and this impaired memory selectivity (Fig. 4a-d, see with alternative inhibitory synaptic plasticity Extended Data Fig. 7j). Furthermore, we specifically blocked CCK^+^ interneurons during memory consolidation in our expanded network model with both CCK^+^ and PV^+^ interneurons and this disrupted memory selectivity (Extended Data Fig. 9l). Note that blocking CCK^+^ interneurons in our model also blocked the plasticity of their efferent synapses (see Methods) in line with several reports that coincident pre- and postsynaptic activity as well as presynaptic activity alone can induce plasticity of *γ*-aminobutyric acid-releasing (GABAergic) synapses onto excitatory neurons.^16^ Interestingly, engram cell turnover was still present in these simulations despite the absence of inhibitory plasticity (Extended Data Fig. 9l). This suggested that excitatory synaptic plasticity alone can drive the emergence of dynamic engrams — consistent with our previous results showing that blocking LTP during memory consolidation leads to engram stabilization (Extended Data Fig. 5a-d). Therefore, our network model predicted that blocking inhibitory synaptic plasticity during memory consolidation prevents the emergence of engram selectivity. To test this prediction, we used a chemogenetic approach to temporarily inhibit DG CCK^+^ interneurons right after fear training (Fig. 4e and Extended Data Fig. 8a right panel, see Methods). In turn, this also blocked the plasticity of CCK^+^ synapses given the known active role of presynaptic activity in the plasticity of GABAergic connections as discussed previously.^16^ Once the effects of the chemogenetic manipulation had subsided, we conducted recall tests and observed that mice in the CCK^+^ inhibition group failed to discriminate between the training and neutral contexts while those in the control group exhibited memory selectivity (Fig. 4f and Extended Data Fig. 8c, compare to Fig 2b-c). Thus, our experimental results supported our model’s prediction that inhibitory synaptic plasticity during memory consolidation is necessary for the emergence of selective engrams.

**Fig. 4.**
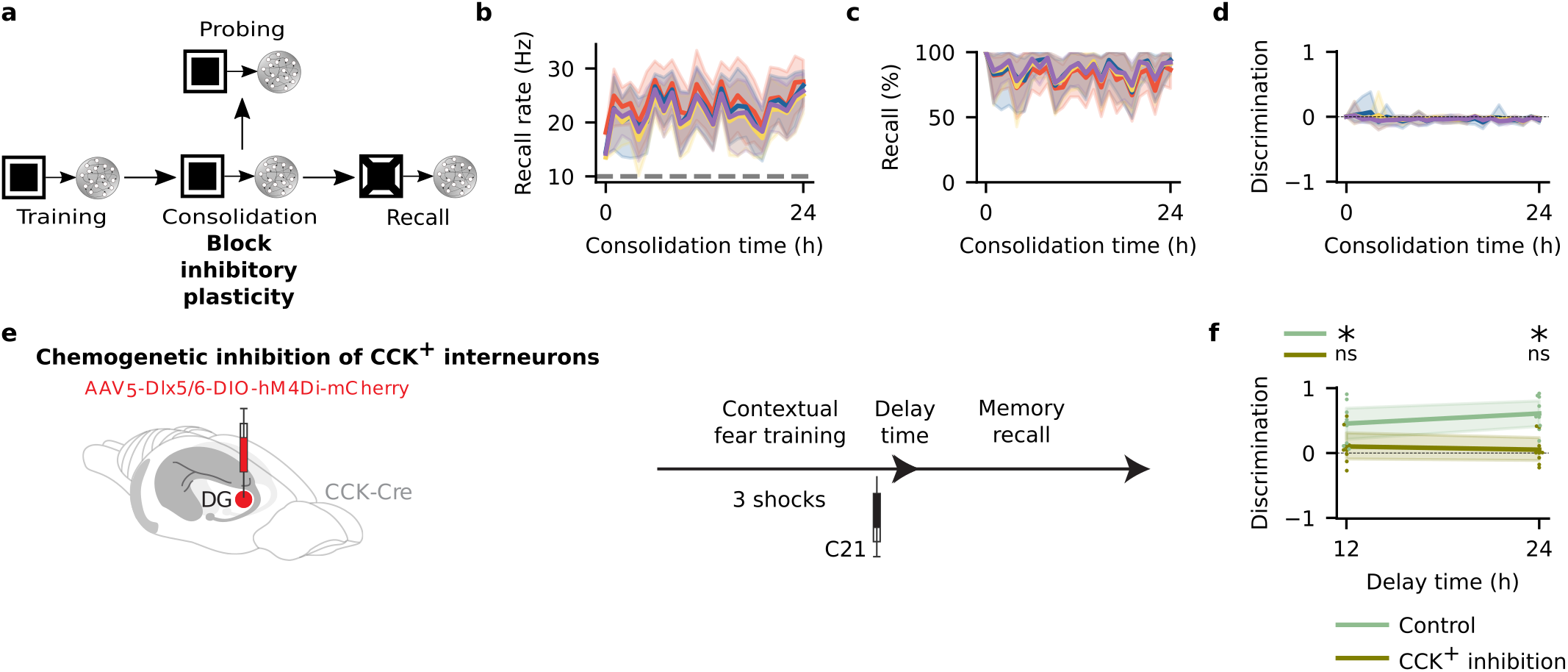
Inhibitory plasticity during memory consolidation carves selective engrams. **a**, Schematic of simulation protocol with blockage of inhibitory synaptic plasticity during consolidation. **b-d**, Means and 99% confidence intervals shown. *n* = 10. **b**, Firing rate of engram cells averaged across all cue presentations during recall in **a** as a function of consolidation time (see Methods). Dashed line indicates threshold *ζ^thr^* = 10 Hz for engram cell activation. Color denotes stimulus as in Fig. 1c. **c**, Memory recall in **a** as a function of consolidation time. Color denotes stimulus as in Fig. 1c. **d**, Discrimination index between recall evoked by cues of the training stimulus and individual novel stimuli in **a** as a function of consolidation time (see Methods). Color denotes stimulus as in Fig. 1c. **e**, Schematic of experimental protocol with chemogenetic inhibition of DG CCK^+^ interneurons. **f**, Discrimination index between freezing levels in the training and neutral contexts in **e** as a function of delay time (see Methods). Means and 99% confidence intervals shown. Two-sided one-sample Wilcoxon signed-rank test against discrimination = 0. Control (*n* = 9 mice per group): for delay = 12 h, *W* = 0.0, *P* = 0.003906; and for delay = 24 h, *W* = 0.0, *P* = 0.003906. CCK^+^ inhibition (*n* = 9 mice per group): for delay = 12 h, *W* = 14.0, *P* = 0.359375; and for delay = 24 h, *W* = 17.0, *P* = 0.570313. *: *P* < 0.05; ns: not significant.

## Discussion

We have shown that memories are encoded by dynamic engrams that become selective with memory consolidation. Previous experiments examined the long-term temporal evolution of neuronal ensembles encoding fear memories in mice prefrontal cortex (PFC) and reported that neurons activated during later recall sessions (14 days after fear training) are more robustly reactivated during remote recall (28 days after fear training) than neurons activated during fear training or earlier recall sessions (1 day after training).^17^ In addition, systems consolidation of fear memories in mice was found to involve training-activated PFC engram cells transitioning from an initial silent (i.e., cannot be reactivated by partial cues) to an active (i.e., can be reactivated by partial cues) state over roughly 12 days while training-activated DG engram cells switch from active to silent in the same timescale.^18^ Moreover, a recent study reported that neurons in the lateral amygdala activated during an initial fear training session become dispensable for memory retrieval after retraining although their artificial reactivation still elicits recall.^19^ Our results extended these findings in a number of ways. First, we showed that DG engrams change in a much shorter timescale, corresponding to hours, with neurons being both removed from and added to the engram cell ensemble. Consequently, this rapid turnover of DG engram cells happens before their active-to-silent transition linked to systems consolidation and without retraining. Second, we proposed a computational framework that identified ongoing synaptic plasticity during memory consolidation as a fundamental mechanism driving changes in engram composition. Third, we found that engram cell turnover is associated with the emergence of selective engrams. Fourth, we showed that inhibition and inhibitory synaptic plasticity are required for expressing and developing memory selectivity, respectively. Lastly, our computational model was able to reconcile the fact that training-activated engram cells are necessary and sufficient for memory recall^4, 5^ with a continuously-changing engram.

Neural representations in the olfactory, visual, parietal, and motor cortices as well as in the hippocampus have been shown to change or drift over time when animals are repeatedly exposed to the same perceptual, navigational, or sensorimotor task.^20–24^ This representational drift was observed in timescales ranging from minutes to weeks depending on the brain region and experimental paradigm. Although our work specifically explored changes in engrams within hours of memory acquisition, our proposition that ongoing post-encoding synaptic plasticity drives continuous changes in neural representations may be a general unifying mechanism that could account for representational drift across regions and tasks. We also associated engram cell turnover to the emergence of memory selectivity. Therefore, our results suggest that representational drift may not necessarily be just a byproduct of synaptic plasticity but effectively have computational and behavioral functions — in line with a proposed role of neural ensemble fluidity in memory updating and flexibility.^25^

Evidence from various pharmacological manipulation studies implies that memory encoding and consolidation is linked to moderate GABAergic transmission.^26^ In particular, administering drugs that enhance GABA transmission either before or after fear training disrupts memory formation. Our work complements these observations by showing that inhibitory neurons become indispensable for memory selectivity via inhibitory synaptic plasticity during consolidation. Taken together, these findings suggest that inhibitory activity levels need to be maintained within a defined range to simultaneously enable the formation of memories and the emergence of selectivity. This is consistent with previous studies reporting that homeostatic inhibitory synaptic plasticity mechanisms can both control firing activity and support functional neuronal networks.^10, 11, 27^ Importantly, we identified CCK^+^ interneurons as mediators of memory selectivity but other interneuron types have also been implicated in the formation and evolution of engrams. For instance, PV^+^ interneurons control the size of engrams in the lateral amygdala^28^ whereas somatostatin-expressing (SST^+^) interneurons constrain the size of hippocampal DG engrams.^29^ Also, CA1 PV^+^ interneuron activity following encoding is required for memory consolidation^30^ while PFC SST interneuron activation and plasticity is critical for memory acquisition and expression.^31^

Past memory models conforming to standard theories of stable memory traces have shed light on several features of network dynamics supporting memory formation and recall.^10, 27, 32^ Our work built on these results to uncover the dynamic nature of memory engrams and the interplay between engram state and memory expression mediated by ongoing synaptic plasticity, possibly offering new directions to investigate pathological conditions characterized by persistently unselective aversive memories such as post-traumatic stress and panic disorders.

## Methods

### Neuron model

We used leaky integrate-and-fire neurons with spike frequency adaption. The membrane voltage *U_i_* of neuron *i* followed:^10^

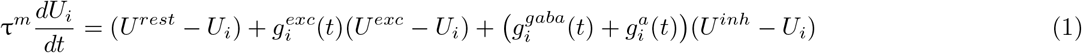

where τ^*m*^ indicates the membrane time constant, *U^rest^* indicates the membrane resting potential, *U^exc^* indicates the excitatory reversal potential, and *U^inh^* indicates the inhibitory reversal potential. The synaptic conductance terms 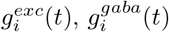, and 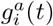 are discussed in the next section.

When the membrane voltage of a neuron *i* exceeds a threshold *ϑ_i_* then it fires a spike. Immediately after firing a spike, the membrane voltage of the neuron is set to 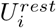 while its firing threshold is temporarily raised to *ϑ^spike^*. In the absence of further spikes, the firing threshold decays to its resting value *ϑ^rest^* with time constant τ^*thr*^ according to:

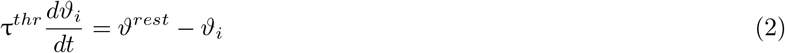

### Synapse model

We employed a conductance-based synaptic input model. The inhibitory synaptic input 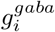 and spike-triggered adaption 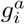 evolved following:^10^

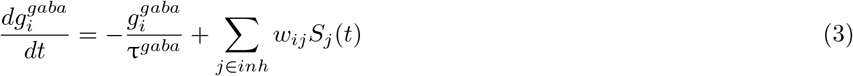

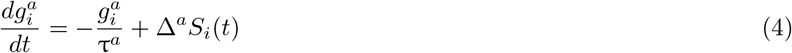

where 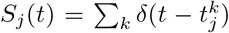 denotes the presynaptic spike train and 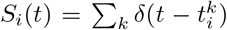 denotes the postsynaptic spike train. In both instances, *δ* indicates the Dirac delta function and 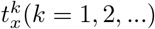 indicates the firing times of neuron *x*. *w_ij_* indicates the weight from neuron *j* to neuron *i*. Δ^*a*^ indicates a fixed adaptation strength. τ^*gaba*^ indicates the GABA decay time constant and τ^*a*^ indicates the adaptation time constant.

Excitatory synaptic input is set by a combination of a fast AMPA-like conductance 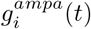 and a slow NMDA-like conductance 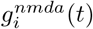:

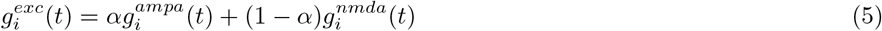

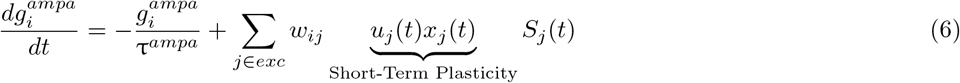

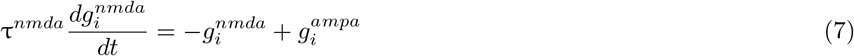

where *α* indicates a constant that determines the relative contribution of 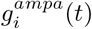 and 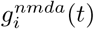 while τ^*ampa*^ and τ^*nmda*^ indicate their respective time constants. *u_j_*(*t*) and *x_j_*(*t*) denote variables that represent the state of short-term plasticity as discussed in the next section.

### Synaptic plasticity model

We designed our synaptic plasticity model after previous work that demonstrated that a combination of Hebbian (i.e., triplet STDP) and non-Hebbian (i.e., heterosynaptic and transmitter-induced) forms of excitatory plasticity is able to support stable memory formation and recall in spiking recurrent neural networks.^10^

#### Short-term plasticity

The variables *u_j_*(*t*) and *x_j_*(*t*) representing the state of short-term plasticity followed:^10^

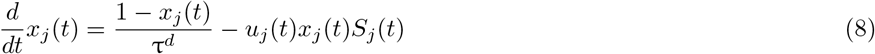

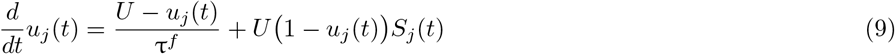

where τ^*d*^ and τ^*f*^ indicate the depression and facilitation time constants, respectively. The parameter *U* indicates the initial release probability.

#### Long-term excitatory synaptic plasticity

Long-term excitatory synaptic plasticity combined triplet STDP,^7^ heterosynaptic plasticity,^8^ and transmitter-induced plasticity^9^ by having an excitatory synaptic weight *w_ij_* from an excitatory neuron *j* to another excitatory neuron *i* evolve according to:^10^

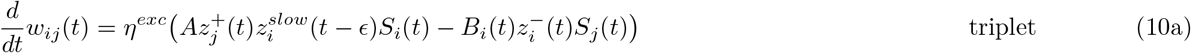

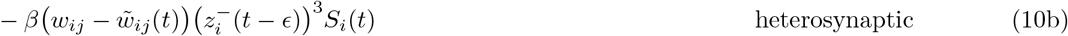

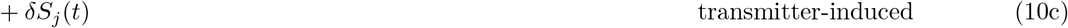

where *η^exc^* (excitatory learning rate), *A* (LTP rate), *β* (heterosynaptic plasticity strength), and *δ* (transmitter-induced plasticity strength) indicate fixed parameters. *ϵ* indicates an infinitesimal offset whose purpose is to ensure that the current action potential is disregarded in the trace. State variables 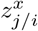 indicate either pre- or postsynaptic traces and each has independent temporal dynamics with time constant τ^*x*^:

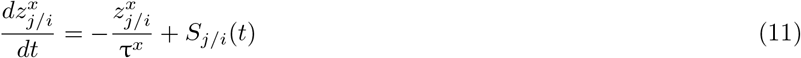

The reference weights 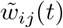 also display independent synaptic consolidation dynamics following the negative gradient of a double-well potential:^10^

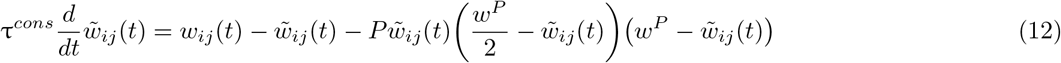

where *P* and *w^P^* indicate fixed parameters and τ^*cons*^ indicates the synaptic consolidation time constant. *P* sets the magnitude of the double-well potential. When *w^P^* = 0.5, an upper stable fixed point is located at 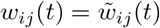 while a lower stable fixed point is located at 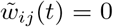. If 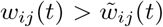 by a small margin, then the upper and lower stable fixed points of 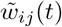 are increased. If 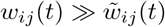, then 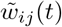 only maintains a single fixed point that has a high value. This synaptic consolidation model is consistent with previous theoretical work^33^ and with synaptic tagging experiments that found that long-lasting LTP relies on events taking place during as well as prior to its initial induction.^34^ In addition, this model assumes the existence of molecular mechanisms that enable synapses to maintain a stable efficacy (i.e., weight) even in the presence of intermittent fluctuations triggered by various factors (e.g., molecular turnover).^35, 36^ Furthermore, the LTD rate *B_i_*(*t*) is subject to homeostatic regulation following:

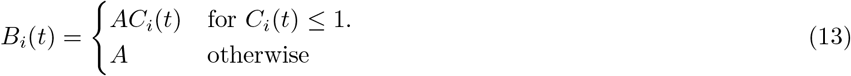

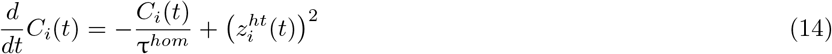

where τ^*hom*^ denotes a time constant and 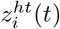 denotes a synaptic trace that follows Equation 11 with its own time constant τ^*ht*^. Finally, excitatory weights were restricted to a range defined by lower and upper bounds 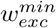 and 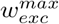, respectively. Nevertheless, excitatory weights never reached their upper bound in our simulations except in some protocols with blockage of neurons or plasticity mechanisms.

#### Inhibitory synaptic plasticity

Inhibitory synaptic plasticity took the form of a network activity-based STDP rule with an inhibitory synaptic weight *w_ij_* from an inhibitory neuron *j* to an excitatory neuron *i* following:^10^

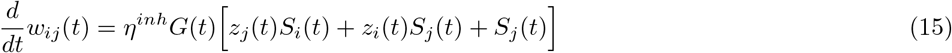

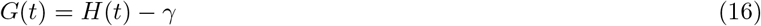

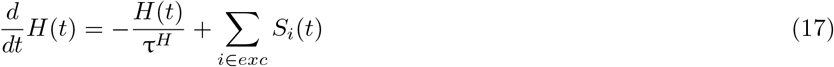

where *η^inh^* indicates a constant inhibitory learning rate and *z_j/i_* indicates either pre- or postsynpatic traces that follow Equation 11 with a shared time constant τ^*iSTDP*^. *G*(*t*) indicates a quantity that linearly depends on the difference between a hypothetical global secreted factor *H*(*t*) and the target local network activity level *γ*. *H*(*t*) indicates a low-pass-filtered version of the spikes fired by the population of excitatory neurons in the local network with time constant τ^*H*^. Note that when the network activity level falls below the target *γ*, *G*(*t*) < 0 and the inhibitory STDP rule (Equation 15) becomes a unidirectional “depression-only” rule.^10^ However, when the network activity level raises above *γ*, the inhibitory rule becomes Hebbian. Thus, the primary goal of inhibitory synaptic plasticity was to control network activity and thereby support the stabilization of the overall network dynamics similar to previous theoretical models.^10, 11, 27^ In Extended Data Fig. 7, we also examined the ability of an alternative version of the inhibitory STDP rule in Equation 15 to support stable network dynamics:

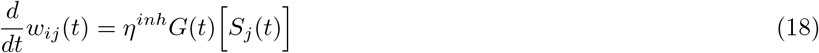

where we retained the presynaptic-only term *S_j_*(*t*) and removed the Hebbian terms *z_j_*(*t*)*S_i_*(*t*) and *z_i_*(*t*)*S_j_*(*t*). In this alternative formulation of the inhibitory rule, *G*(*t*) and *H*(*t*) also follow Equations 16 and 17, respectively. Lastly, inhibitory weights were restricted to the range between 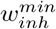 and 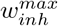 but they never reached their upper limit in our simulations except in some protocols with blockage of neurons or plasticity mechanisms.

### Network model

The model consisted of a stimulus population of *N_stim_* = 4, 096 Poisson neurons and a recurrent neural network corresponding to the hippocampus. Importantly, the following anatomical evidence has motivated the use of a recurrent neural network to model the hippocampus: recurrent excitatory synapses in CA3^37^ and to a less extent in CA1;^38^ feedforward synapses from dentate gyrus to CA3 and from CA3 to CA1;^39^ “backprojection” synapses from CA1 to CA3 and dentate gyrus^40^ and from CA3 to dentate gyrus;^41^ and recurrent inhibitory synapses in CA1,^40^ CA3,^37^ and dentate gyrus.^42^ The hippocampus network was composed of *N_exc_* = 4, 096 excitatory neurons and *N_inh_* = 1, 024 inhibitory neurons that were recurrently connected. Recurrent excitatory synapses onto excitatory neurons exhibited short- and long-term excitatory synaptic plasticity whereas excitatory synapses onto inhibitory neurons displayed only short-term plasticity. Inhibitory synapses onto inhibitory neurons were static whereas inhibitory synapses onto excitatory neurons exhibited inhibitory synaptic plasticity. Feedforward excitatory synapses from the stimulus population to excitatory neurons in the hippocampus exhibited short- and long-term excitatory synaptic plasticity. Recurrent synapses were randomly initialized following a uniform distribution while feedforward synapses from the stimulus population had circular receptive fields centered at random locations (i.e., each excitatory neuron in the hippocampus received projections from a small circular area in the stimulus population of radius *R_hpc_* whose random center location followed a uniform distribution). Plasticity mechanisms were active in the entirety of all simulations except in protocols where plasticity was purposefully blocked. Recurrent synapses had a connection probability *ϵ_rec_* and were initialized with specific weights (i.e., *w^EE^*, *w^EI^*, *w^II^*, and *w^IE^*). Feedforward synapses had an initial weight *w_stim_*. For a complete list of network parameters, see Supplementary Table 1.

### Network simulation

Network simulations comprised multiple phases: burn-in, training, consolidation, probing, and recall. Each simulation began with a brief burn-in period of duration *T_burn_* that stabilized the activity of the hippocampus network under background input from the stimulus population at rate *ν^bg^*. Next, the training stimulus (i.e., Fig. 1c) was randomly presented to the hippocampus network in the training phase of duration *T_training_* with stimulus-off and stimulus-on intervals both drawn from exponential distributions with means 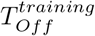 and 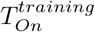, respectively. Following the training phase, the training stimulus was randomly reactivated in the consolidation phase with stimulus-off and stimulus-on periods also drawn from exponential distributions but with means 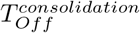 and 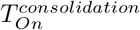, respectively. This was motivated by recent experiments that showed that training-activated sensory neurons are selectively reactivated during post-training sleep and that this reactivation is essential for memory selectivity.^12^ Hence, our model aimed to capture the role of sleep-dependent sensory reactivation in memory consolidation. At regular intervals throughout the consolidation phase (i.e., consolidation time = 0, 1, …, *T_consolidation_* hours), the network advanced to the probing phase when the training stimulus was randomly presented for a brief period *T_probing_* with stimulus-off and stimulus-on intervals drawn from exponential distributions with means 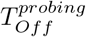 and 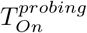, respectively. This allowed us to identify the set of engram cells that encode the training stimulus at any consolidation time point (see next section). We also subjected the network to the recall phase of duration *T_recall_* after each consolidation interval (0, 1, …, *T_consolidation_* hours). For each stimulus (i.e., either the training stimulus or one of three novel stimuli, see Fig. 1c), a separate recall session was conducted with partial cues of that individual stimulus being presented to the network. Hence, there was a total of four distinct parallel recall sessions (i.e., one for the training stimulus and three for the novel stimuli) for each sampled consolidation interval. Cue-off and cue-on intervals also followed exponential distributions with means 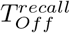 and 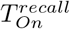, respectively. Each stimulus and partial cue was depicted in a 64 x 64 grid in Fig. 1c and the overlap between the training stimulus and each novel stimulus was shown in Extended Data Fig. 1a. During presentation or reactivation of a stimulus or cue, the stimulus population kept firing at the background rate *ν^bg^* but the stimulus neurons that corresponded to a given stimulus or cue selectively increased their firing rate to *ν^stim^*. When blocking neurons (i.e., stimulus, excitatory, or inhibitory neurons), their output and all their efferent synapses were blocked. For a complete list of simulation parameters, see Supplementary Table 1.

### Engram cells and recall metrics in simulations

In our simulations, engram cells were identified by computing the average stimulus-evoked firing rate of each neuron in the hippocampus network at different time points. First, a neuron was said to be an engram cell encoding the training stimulus at the end of the training phase if its average stimulus-evoked firing rate was above the threshold *ζ^thr^* = 10 Hz for the last Δ*t^eng^* = 60 seconds of the training phase. Post-training, we used the probing phase to identify engram cells after consolidation time had elapsed. Specifically, following consolidation time = 0, 1, …, *T_consolidation_* hours, a neuron was regarded an engram cell encoding the training stimulus if its average stimulus-evoked firing rate was above the threshold *ζ^thr^* = 10 Hz during the entire probing phase of duration *T_probing_* = 60 seconds. Consequently, a neuron may be an engram cell at one point and subsequently no longer be an engram cell at a later time point (i.e., the neuron “dropped out of” the engram^2^). Conversely, a neuron may not be an engram cell initially but later may become an engram cell (i.e., the neuron “dropped into” the engram^2^). In addition, a neuron may alternate between being and not being an engram cell over the course of consolidation. Naturally, a neuron may be an engram cell or a non-engram cell at the end of the training phase and remain so throughout the consolidation period. To examine the evolution of engrams with memory consolidation in our simulations, we measured the overlap between engram cells identified in the probing phase (i.e., probing-activated or probing^+^ engram cells) and in the training phase (i.e., training-activated or training^+^ engram cells) as a fraction of training^+^, probing^+^, or all neurons in the hippocampus network. We also tracked the size of the probing^+^ engram cell ensemble as fraction of all neurons in the network as well as the hour-to-hour overlap between probing^+^ engram cell ensembles. Moreover, training stimulus neurons were considered stable sensory engram cells given the prominent role of training-activated sensory neurons in memory storage and consolidation.^12^ Furthermore, an engram cell ensemble at a given time point was taken as activated upon presentation of a partial cue of either the training or a novel stimulus if its population firing rate (i.e., average firing rate computed over all engram cells in the ensemble) was above the threshold *ζ^thr^* = 10 Hz during presentation of the cue. We then measured recall as the number of instances when the engram cell ensemble was activated following cue presentation divided by the total number of cue presentations in the recall phase. Moreover, we defined recall rate as the population firing rate of engram cells evoked by partial cues averaged across all cue presentations in the recall phase. We computed separate recall metrics for the training and novel stimuli given that cues of a single stimulus were presented in a given recall session (see previous section).

### Mice

C57BL/6J wild type male mice were obtained from Jackson Laboratory. Experiments using CCK-Cre mice employed the CCK-IRES-Cre knock-in line (Stock No. 012706, Jackson Laboratory). Knock-in mice were maintained as hemizygotes. Mice had access to food and water *ad libitum* and were socially housed in numbers of two to five littermates until surgery. Following surgery, mice were single housed. For behavioral experiments, all mice were male and 2-4 months old. All experiments were conducted in accordance with U.S. National Institutes of Health (NIH) guidelines and the Massachusetts Institute of Technology Department of Comparative Medicine and Committee on Animal Care.

### Mouse behavior

Experiments were conducted during the light cycle (7 am to 7 pm). Mice were randomly assigned to experimental groups for specific behavioral assays immediately after surgery. Mice were habituated to investigator handling for 1-2 minutes on three consecutive days. Handling took place in the holding room where the mice were housed. Prior to each handling session, mice were transported by wheeled cart to and from the vicinity of the behavior rooms to habituate them to the journey. All behavior experiments were analyzed blind to experimental group. Following behavioral protocols, brain sections were prepared to confirm efficient viral labeling in hippocampal DG. Animals lacking adequate labeling were excluded prior to behavior quantification.

#### Contextual fear conditioning

Two distinct contexts were employed. The conditioning context was a 29 × 25 × 22 cm chamber with grid floors, dim white lighting, and scented with 0.25% benzaldehyde. The neutral context consisted of a 29 × 25 × 22 cm chamber with white perspex floors, red lighting, and scented with 1% acetic acid. All mice were conditioned (120 s exploration, one 0.65 mA shock of 2 s duration at 120 s, 60 s post-shock period, second 0.65 mA shock of 2 s duration at 180 s, 60 s post-shock period, third 0.65 mA shock of 2 s duration at 240 s, 60 s post-shock period). Following different intervals (1, 5, 12, 18, 24 h), mice were tested in the neutral context (3 min) followed by a recall test in the conditioning context (3 min) ∼1 hour later. Floors of chambers were cleaned with quatricide before and between runs. Mice were transported to and from the experimental room in their home cages using a wheeled cart. For experiments that included optogenetic manipulations (including Cal-Light active neuron labeling), the behavior chamber ceilings were customized to hold a rotary joint (Doric Lenses) connected to two 0.3 m optic fibers. All mice had optic fibers attached to their optic fiber implants prior to training and recall tests. Since optogenetic manipulations (i.e., optic fibers) interfered with automated motion detection, freezing behavior was manually quantified for these experiments.

### Surgery

Animals were anesthetized with isoflurane for stereotaxic injections and were given 1 mg kg^−1^ meloxicam as analgesic prior to incisions. Virus was injected at 70 nl min^−1^ using a glass micropipette attached to a 10 ml Hamilton microsyringe. The needle was lowered to the target site and remained for 5 min before beginning the injection. After the injection, the needle stayed for 10 min before it was withdrawn. For labeling and behavioral manipulation experiments using optogenetics, implants were lowered right above injection sites. The implant was secured to the skull with two jewelry screws, adhesive cement (C&B Metabond), and dental cement. Mice were given 1-2 mg kg^−1^ sustained-release buprenorphine as analgesic after surgeries and allowed to recover for at least 2 weeks before behavioral experiments.

### Activity-dependent labeling

For Cal-Light experiments, a cocktail of three viruses was injected into hippocampal DG of wild type mice followed by bilateral optic fiber implants (200 μm core diameter, Newdoon). Viruses were AAV_5_-TM-CaM-NES-TEV-N-AsLOV2-TEVseq-tTA (Addgene plasmid #92392), AAV_5_-M13-TEV-C-P2A-tdTomato (Addgene plasmid #92391), and AAV_5_-TRE-EGFP (Addgene plasmid #89875). These constructs were serotyped with AAV_5_ coat proteins and packaged by the Viral Core at Boston Children’s Hospital (∼4 × 1012 GC ml^−1^ viral titer). Coordinates for hippocampal DG were AP −2.0 mm, +/− 1.0 mm ML, −2.1 mm DV, and 250 nl of the virus cocktail was injected per hemisphere. Following recovery, active neurons during fear conditioning were tagged with EGFP by delivery of continuous blue light (473 nm, 15 mW at patch cord tip) for the entire duration of the training session (∼5 min). For *c-Fos* staining, animals were perfused 60 min after the specific behavioral epoch.

### Immunohistochemistry

Mice were dispatched using an overdose of isoflurane and transcardially perfused with PBS, followed by 4% paraformaldehyde (PFA). Brains were extracted and incubated in 4% PFA at room temperature overnight. Brains were transferred to PBS and 50 μm coronal slices were prepared using a vibratome. For immunostaining, each slice was placed in PBS + 0.2% Triton X-100 (PBS-T), with 5% normal goat serum for 1 h and then incubated with primary antibody at 4°C for 24 h. Slices then underwent three wash steps for 10 min each in PBS-T, followed by a 2 h incubation with secondary antibody. After three more wash steps of 10 min each in PBS-T, slices were mounted on microscope slides. Antibodies used for staining were as follows: chicken anti-GFP (1:1000, Life Technologies) and anti-chicken Alexa-488 (1:1000), rabbit anti-cFos (1:500, Cell Signaling Technology) and anti-rabbit Alexa-555 (1:300), and nuclei were stained with DAPI (1:3000, Sigma).

### Cell counting

Brain sections were imaged with a 10X magnification objective on an Olympus fluorescent microscope. Images were processed using ImageJ, and quantifications were performed manually from 3-5 sections per animal. All counting experiments were conducted blind to experimental group. Researcher 1 trained the animals, prepared slices, and randomized images, while Researcher 2 performed cell counting. Neuronal cell counts were normalized to the number of DAPI^+^ cells in the field of view.

### Chemogenetic and optogenetic manipulations

For chemogenetic (i.e., hM4Di) neuronal activity manipulation experiments, we generated and injected an AAV_5_-Dlx5/6-DIO-hM4Di-mCherry virus (∼5.3 × 1012 GC ml^−1^ viral titer) into hippocampal DG of CCK-Cre mice. We used the second-generation DREADD agonist known as compound 21 (C21). This agonist was purchased in a water-soluble dihydrochloride form (Hello Bio). For each mouse, optimal chemogenetic activity was achieved using a target concentration of 2 mg kg^−1^ (injected intraperitoneally), which in this study was injected right after fear conditioning to inhibit DG CCK^+^ interneurons during the subsequent cellular consolidation window. For optogenetic neuronal activity manipulation experiments, we generated and injected an AAV_5_-Dlx5/6-DIO-eArch3.0-eYFP virus (∼3.8 × 1012 GC ml^−1^ viral titer) into hippocampal DG of CCK-Cre mice, followed by bilateral optic fiber implants targeting hippocampal DG. eArch3.0 was activated with a 570 nm laser (10 mW, constant green light).

### Discrimination index

We defined a memory recall discrimination index for our network simulations as:

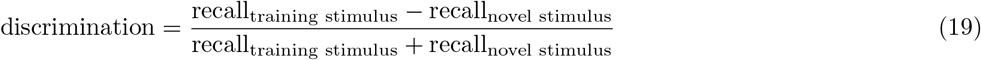

where recall_training stimulus_ and recall_novel stimulus_ denote recall levels when cues of the training stimulus and a novel stimulus are presented in the recall phase of our simulation protocol, respectively. Note that a separate discrimination index was calculated for each novel stimulus. We also defined an analogous memory recall discrimination index for our contextual fear conditioning experiments:

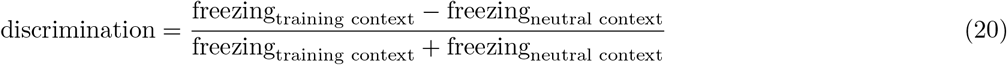

where freezing_training context_ and freezing_neutral context_ denote freezing levels during memory recall in the training and neutral contexts of our experimental protocol, respectively. In both cases, the discrimination index measured the ability of the network model (Equation 19) or mice (Equation 20) to selectively recall the encoded memory only when cues of the training experience were presented.

### Statistics

Freezing levels in experiments were compared at multiple time points using a two-sided Wilcoxon signed-rank test between freezing in the training and neutral contexts. Each memory discrimination index between freezing levels in the training and neutral contexts in experiments was compared against an average discrimination = 0 using a two-sided one-sample Wilcoxon signed-rank test. Memory recall at consolidation time = 24 h in the control simulation (Fig. 1f) and in the simulation with blockage of LTP during consolidation (Extended Data Fig. 5f) were compared using a two-sided Wilcoxon signed-rank test. Quantities compared using a Wilcoxon signed-rank test met required assumptions of this test (i.e., dependent samples, independence of paired observations, continuous dependent variable, and ordinal level of measurement). A one-way ANOVA followed by Tukey’s HSD post hoc test was used to compare ensemble overlap between neurons activated during both recall and training in experiments (*c-Fos*^+^ ∩ EGFP^+^) when memory recall was evaluated either in the training context, neutral context, or home cage. *c-Fos*^+^ ∩ EGFP^+^ as a fraction of training-activated neurons (EGFP^+^), recall-activated neurons (*c-Fos*^+^), and neuronal cell count (DAPI^+^) were each tested separately. Ensemble overlap data met required assumptions of one-way ANOVA (i.e., independent samples, normality of residuals, homogeneity of variances, and continuous dependent variable). For each test, the null hypothesis was rejected at the *P* < 0.05 significance level. Sample sizes were chosen based on previous studies.^4, 5, 17^

### Simulation and data analysis details

We employed the forward Euler method to update neuronal state variables with a time step Δ = 0.1 ms (with the exception of reference weights 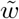 for which we set a longer time step Δ_*long*_ = 1.2 s for efficiency reasons). Population activity (i.e., average firing rate over all neurons in a given ensemble) was computed with a temporal resolution of 10 ms without smoothing or convolution. We computed 99% confidence intervals for mean metrics (i.e., ensemble overlap, recall firing rate, recall, freezing level, and discrimination index) using a non-parametric bootstrap to provide an estimate of uncertainty and to facilitate their visualization.

### Code details

Simulation code was written in C++ employing the Auryn framework for spiking neural network simulation.^43^ After several preliminary simulations, we found that setting the number of Message Passing Interface (MPI) ranks *N_ranks_* = 16 minimized our simulation time with Auryn. Code used to analyze simulation output and experimental data was written in Python 3.8.

## Data availability

The data necessary to reproduce the simulations reported in this study will be made publicly available upon publication. Individual mouse behavioral and cell counting data will also be made available upon publication.

## Code availability

The code used to perform the simulations and data analyses in this work will be made available in a public repository upon publication.

## Acknowledgments

This work was funded by the President’s PhD Scholarship from Imperial College London (D.F.T.), BBSRC (BB/N013956/1 and BB/N019008/1) (C.C.), Wellcome Trust (200790/Z/16/Z) (C.C.), the Simons Foundation (564408) (C.C.), and the EPSRC (EP/R035806/1) (CC). The J. Douglas Tan Postdoctoral Fellowship supported Y.Z. The Warren Alpert Distinguished Scholar Award and the NIH 1K99NS125131-01 supported D.S.R.

## Author contributions

D.F.T. conceptualized the model, performed the simulations, analyzed simulation output and experimental data, and wrote the original draft of the manuscript. Y.Z. and D.S.R. designed the experiments, performed the experiments, and wrote the original draft of the manuscript. T.A. generated AAV constructs used in the experiments. S.S., D.S.R., and C.C. supervised the project. All authors reviewed and edited the manuscript.

## Additional Information

**Supplementary Information** is available for this paper.

---

**Extended Data Fig. 1.**
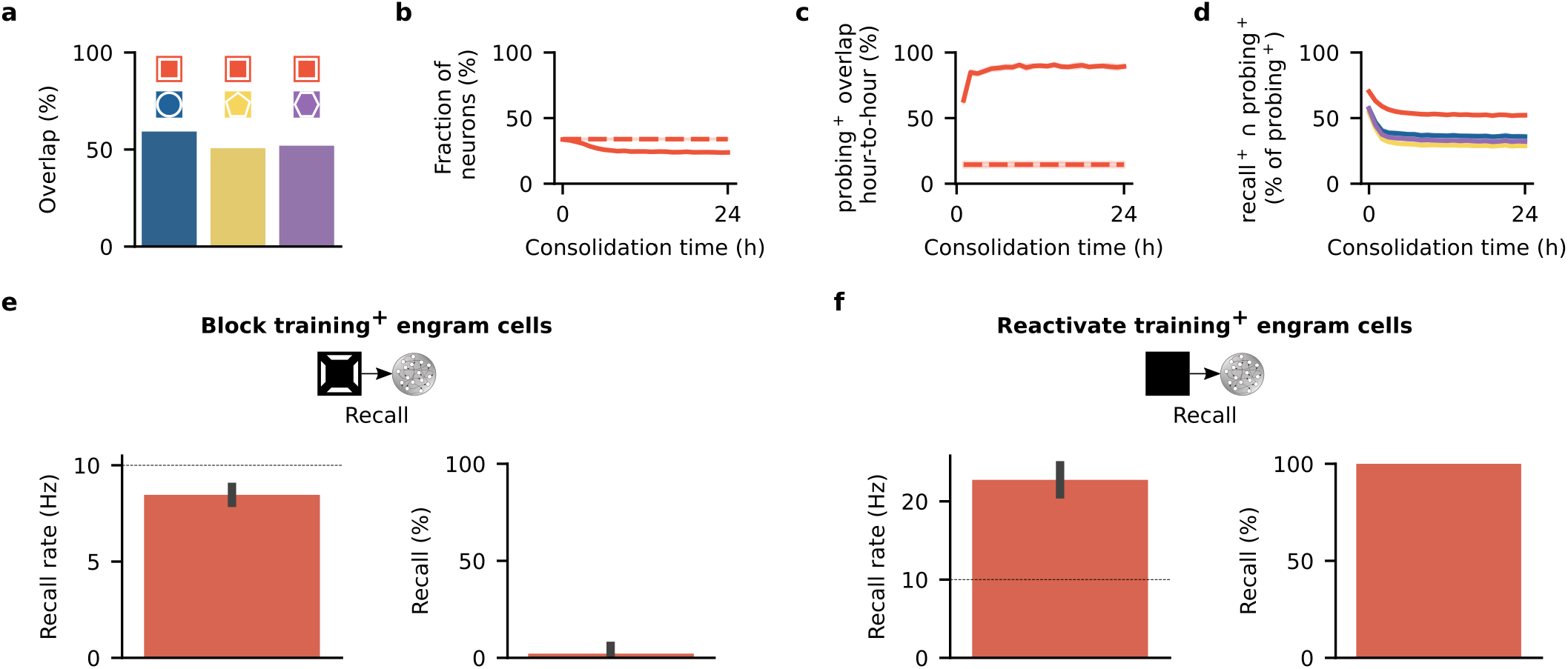
Training-activated engram cells remain necessary and sufficient for memory recall. **a**, Overlap between the training stimulus and each novel stimulus in Fig. 1c as a fraction of training stimulus neurons. **b-d** Analysis of engram cells in Fig. 1b. Means and 99% confidence intervals shown. *n* = 10. **b**, Ensemble of engram cells as of fraction of all neurons (see Methods). Dashed line indicates engram cell ensemble at the end of training. **c**, Ensemble overlap between probing-activated engram cells at consolidation time = *t* and *t*-1 h as a fraction of engram cells at consolidation time = *t*-1 h. Dashed line indicates ensemble of neurons that remained part of the engram in all sampled time points (i.e., consolidation time = 0, 1, …, 24 h) as a fraction of engram cells at consolidation time = 0 h (i.e., training-activated engram cells). **d**, Fraction of probing-activated engram cells reactivated during recall as a function of consolidation time. Color denotes stimulus as in Fig. 1c. **e-f**, Analysis of memory recall when training-activated engram cells in Fig. 1b are manipulated. Means and standard deviations shown. *n* = 10. **e**, Analysis of memory recall after 24 hours of consolidation when training-activated engram cells are blocked during recall with cues of the training stimulus in the protocol in Fig. 1b. Left: firing rate of engram cells averaged across all cue presentations during recall (dashed line indicates threshold *ζ^thr^* = 10 Hz for engram cell activation). Right: memory recall. **f**, Analysis of memory recall after 24 hours of consolidation when training-activated engram cells are artificially reactivated during recall without any stimulus cues in the protocol in Fig. 1b. Left: firing rate of engram cells averaged across all reactivation intervals during recall (dashed line indicates threshold *ζ^thr^* = 10 Hz for engram cell activation). Right: memory recall (note that standard deviation = 0 as recall = 100% in all trials).

**Extended Data Fig. 2.**
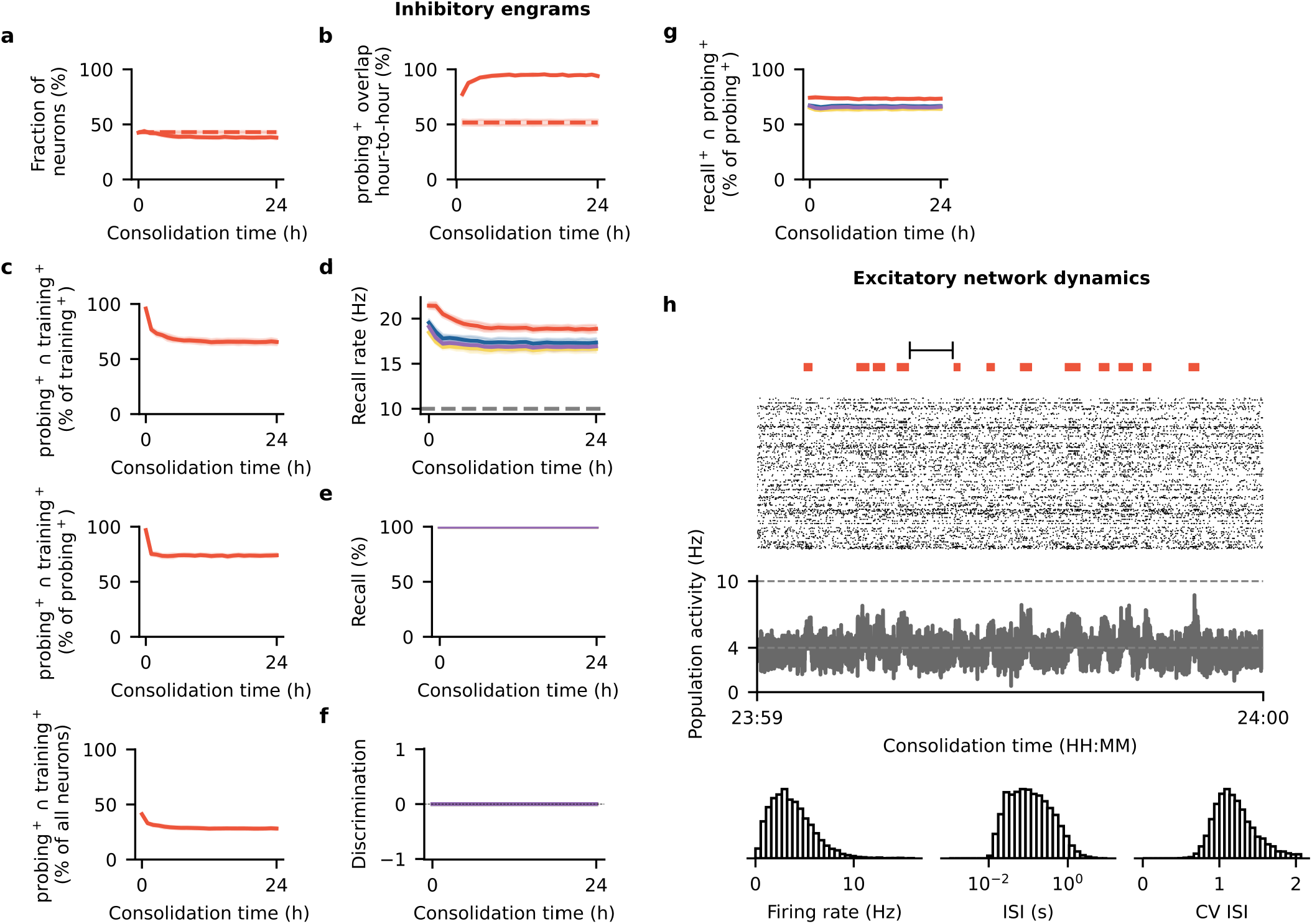
Dynamic and unselective inhibitory engrams control network activity. **a-c**, Post-encoding evolution of inhibitory engram cells in Fig. 1b. Means and 99% confidence intervals shown. *n* = 10. **a**, Ensemble of inhibitory engram cells as of fraction of all inhibitory neurons (see Methods). Dashed line indicates inhibitory engram cell ensemble at the end of training. **b**, Ensemble overlap between probing-activated inhibitory engram cells at consolidation time = *t* and *t*-1 h as a fraction of inhibitory engram cells at consolidation time = *t*-1 h. Dashed line indicates ensemble of inhibitory neurons that remained part of the inhibitory engram in all sampled time points (i.e., consolidation time = 0, 1, …, 24 h) as a fraction of inhibitory engram cells at consolidation time = 0 h (i.e., training-activated inhibitory engram cells). **c**, Ensemble overlap between inhibitory engram cells activated during both probing and training as a fraction of training-activated inhibitory engram cells (top), probing-activated inhibitory engram cells (middle), and all inhibitory neurons in the network (bottom). **d-g**, Analysis of inhibitory engram cells during recall in Fig. 1b. Means and 99% confidence intervals shown. *n* = 10. Color denotes stimulus as in Fig. 1c. **d**, Recall rate of inhibitory engram cells as a function of consolidation time. Dashed line indicates threshold *ζ^thr^* = 10 Hz for engram cell activation. **e**, Recall of inhibitory engram cells as a function of consolidation time. **f**, Discrimination index between recall of inhibitory engram cells evoked by cues of the training stimulus and individual novel stimuli as a function of consolidation time. **g**, Fraction of probing-activated inhibitory engram cells reactivated during recall as a function of consolidation time. **h**, Analysis of excitatory network activity in a a 60-second interval during consolidation in Fig. 1b. From top to bottom: training stimulus reactivation times, spike raster of 256 randomly-chosen excitatory neurons (for clarity, only every fifth spike is shown), population activity (i.e., average firing rate) of all excitatory neurons (dashed lines indicate target activity level *γ* = 4 Hz and threshold *ζ^thr^* = 10 Hz for engram cell activation), and network statistics for the interval marked at the top (from left to right: histograms of firing rates, interspike intervals, and coefficient of variation of interspike intervals). Population activity is shown without smoothing or convolution (see Methods). Representative trial shown.

**Extended Data Fig. 3.**
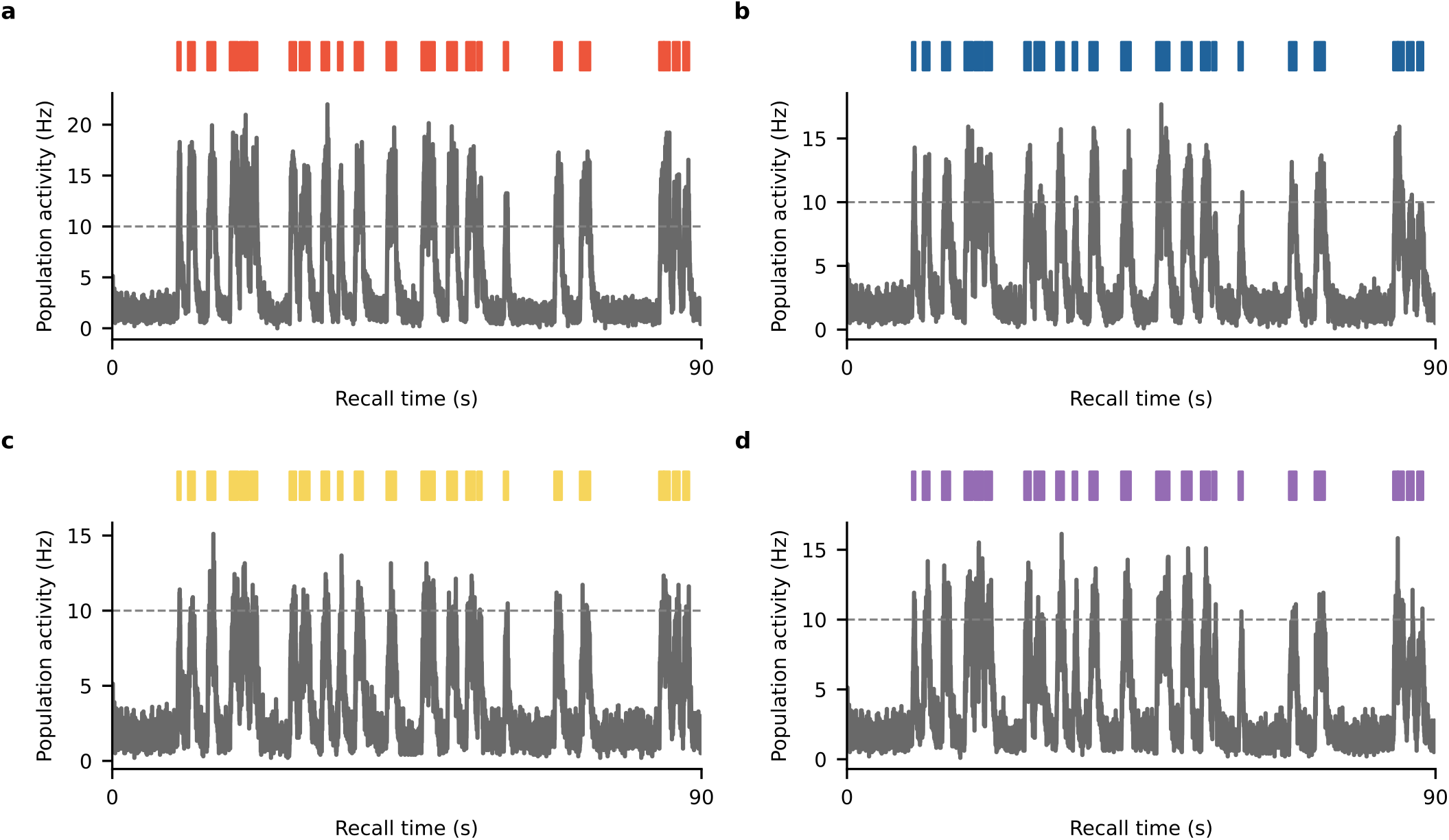
Dynamic engrams are selectively reactivated by cues of the training stimulus. **a-d**, Population activity (i.e., average firing rate) of engram cells during recall after 24 hours of consolidation in Fig. 1b. **a**, Population activity of engram cells when cues of the training stimulus are presented during recall. Population activity is shown without smoothing or convolution (see Methods). Cue presentation times indicated at the top where color denotes stimulus in Fig. 1c. Dashed line indicates threshold *ζ^thr^* = 10 Hz for engram cell activation. When presenting a cue, engram cell ensemble is considered activated if its average firing rate is above the threshold *ζ^thr^* = 10 Hz during the cue presentation interval (see Methods). Representative trial shown. **b**, Same as **a** but when cues of a novel stimulus (i.e., circle in Fig. 1c) are presented. **c**, Same as **a** but when cues of a novel stimulus (i.e., pentagon in Fig. 1c) are presented. **d**, Same as **a** but when cues of a novel stimulus (i.e., hexagon in Fig. 1c) are presented.

**Extended Data Fig. 4.**
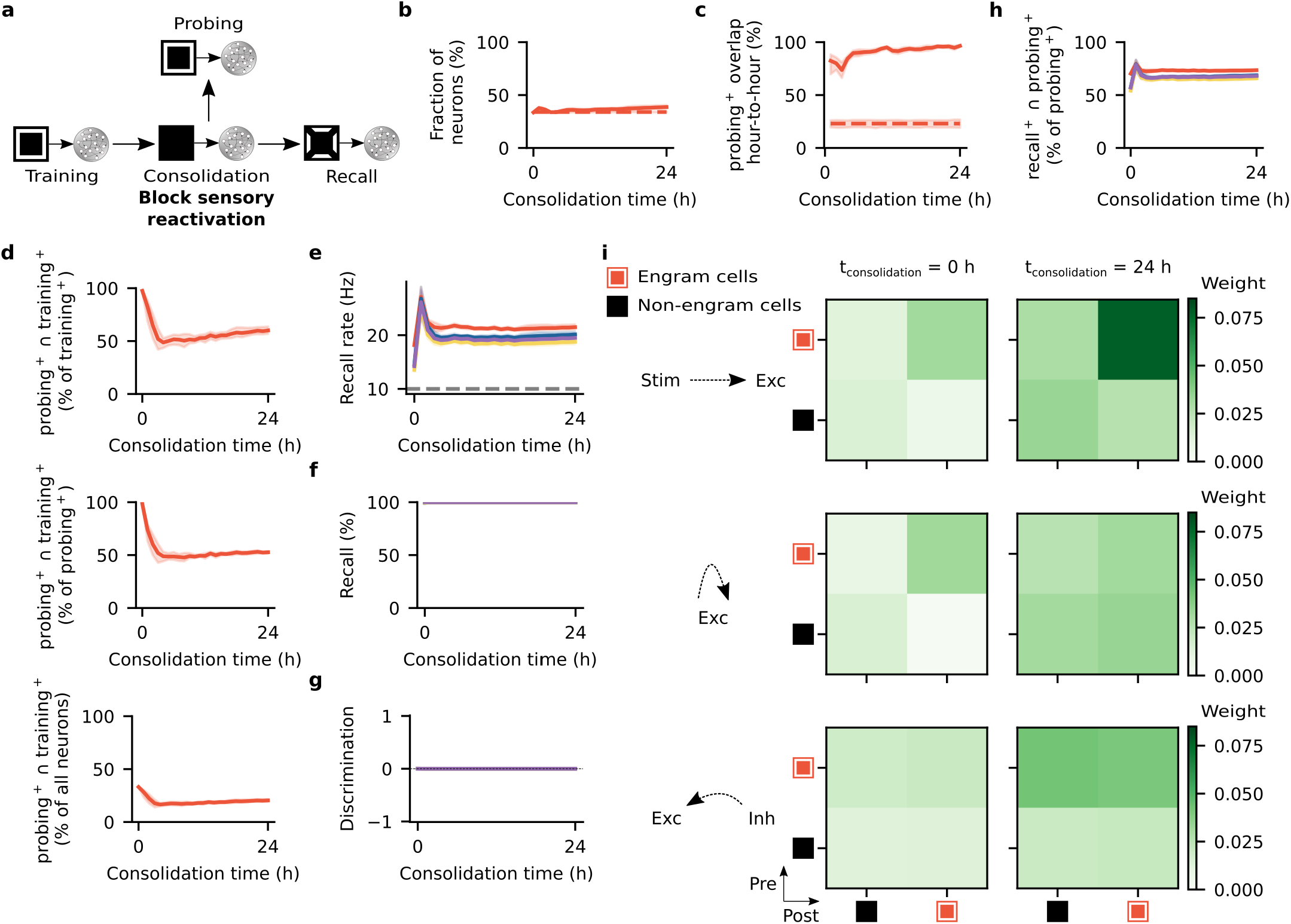
Sensory reactivation during memory consolidation is essential for selectivity. **a**, Schematic of simulation protocol with blockage of training-activated stimulus neurons during consolidation. Network and simulation parameters as in Fig. 1b. **b-d**, Post-encoding evolution of engram cells in **a**. Means and 99% confidence intervals shown. *n* = 10. **b**, Ensemble of engram cells as of fraction of all neurons (see Methods). Dashed line indicates engram cell ensemble at the end of training. **c**, Ensemble overlap between probing-activated engram cells at consolidation time = *t* and *t*-1 h as a fraction of engram cells at consolidation time = *t*-1 h. Dashed line indicates ensemble of neurons that remained part of the engram in all sampled time points (i.e., consolidation time = 0, 1, …, 24 h) as a fraction of engram cells at consolidation time = 0 h (i.e., training-activated engram cells). **d**, Ensemble overlap between engram cells activated during both probing and training as a fraction of training-activated engram cells (top), probing-activated engram cells (middle), and all neurons in the network (bottom). **e-h**, Analysis of memory recall in **a**. Means and 99% confidence intervals shown. *n* = 10. Color denotes stimulus as in Fig. 1c. **e**, Firing rate of engram cells averaged across all cue presentations during recall as a function of consolidation time. Dashed line indicates threshold *ζ^thr^* = 10 Hz for engram cell activation. **f**, Memory recall as a function of consolidation time. **g**, Discrimination index between recall evoked by cues of the training stimulus and individual novel stimuli as a function of consolidation time (see Methods). **h**, Fraction of probing-activated engram cells reactivated during recall as a function of consolidation time. **i**, Mean weight strength of plastic synapses in the network in **a** clustered according to engram cell status (i.e., engram and non-engram cells). Top: feedforward excitatory synapses onto excitatory neurons. Middle: recurrent excitatory synapses onto excitatory neurons. Bottom: recurrent inhibitory synapses onto excitatory neurons. Left: at the end of the training phase. Right: after 24 hours of consolidation. Representative trial shown.

**Extended Data Fig. 5.**
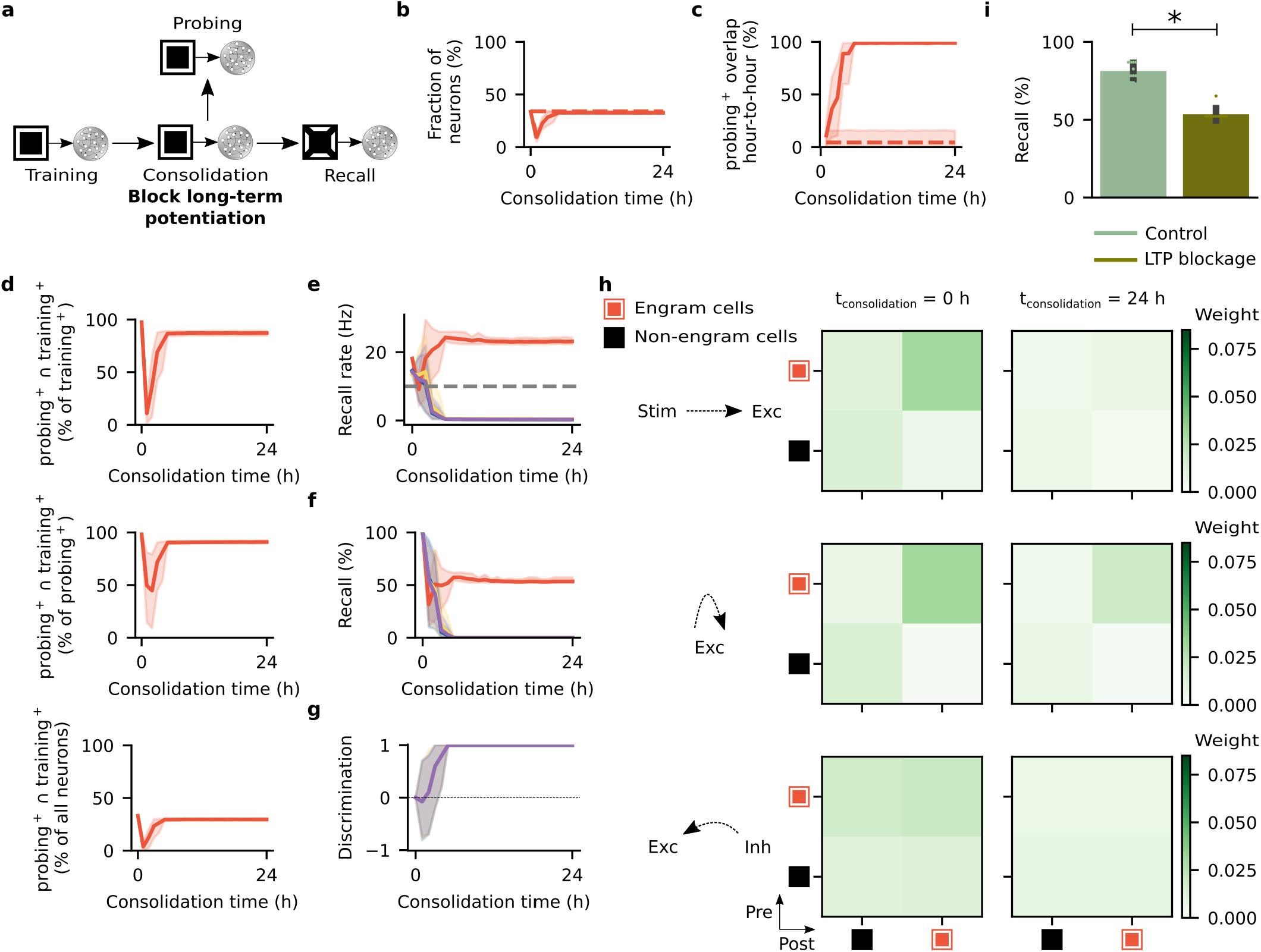
Blocking long-term potentiation during memory consolidation impairs recall. **a**, Schematic of simulation protocol with blockage of long-term potentiation (LTP) during consolidation. LTP induced by triplet STDP and transmitter-induced plasticity are both blocked. Network and simulation parameters as in Fig. 1b except that *A* = *δ* = 0 during consolidation (see Methods). **b-d**, Post-encoding evolution of engram cells in **a**. Means and 99% confidence intervals shown. *n* = 10. **b**, Ensemble of engram cells as of fraction of all neurons. Dashed line indicates engram cell ensemble at the end of training. **c**, Ensemble overlap between probing-activated engram cells at consolidation time = *t* and *t*-1 h as a fraction of engram cells at consolidation time = *t*-1 h. Dashed line indicates ensemble of neurons that remained part of the engram in all sampled time points (i.e., consolidation time = 0, 1, …, 24 h) as a fraction of training-activated engram cells. **d**, Ensemble overlap between engram cells activated during both probing and training as a fraction of training-activated engram cells (top), probing-activated engram cells (middle), and all neurons in the network (bottom). **e-g**, Analysis of memory recall in **a**. Means and 99% confidence intervals shown. *n* = 10. Color denotes stimulus as in Fig. 1c. **e**, Firing rate of engram cells averaged across all cue presentations during recall as a function of consolidation time. Dashed line indicates threshold *ζ^thr^* = 10 Hz for engram cell activation. **f**, Memory recall as a function of consolidation time. **g**, Discrimination index as a function of consolidation time. **h**, Mean weight strength of plastic synapses in the network in **a** clustered according to engram cell status (i.e., engram and non-engram cells). Top: feedforward excitatory synapses onto excitatory neurons. Middle: recurrent excitatory synapses onto excitatory neurons. Bottom: recurrent inhibitory synapses onto excitatory neurons. Left: at the end of the training phase. Right: after 24 hours of consolidation. Representative trial shown. **i**, Memory recall evoked by cues of the training stimulus at t_consolidation_ = 24 h in Fig. 1f (control) and in **f** (LTP blockage) (same data). Two-sided Wilcoxon signed-rank test. *W* = 0.0, *P* = 0.001953. Means and standard deviations shown. *n* = 10. *: *P* < 0.05.

**Extended Data Fig. 6.**
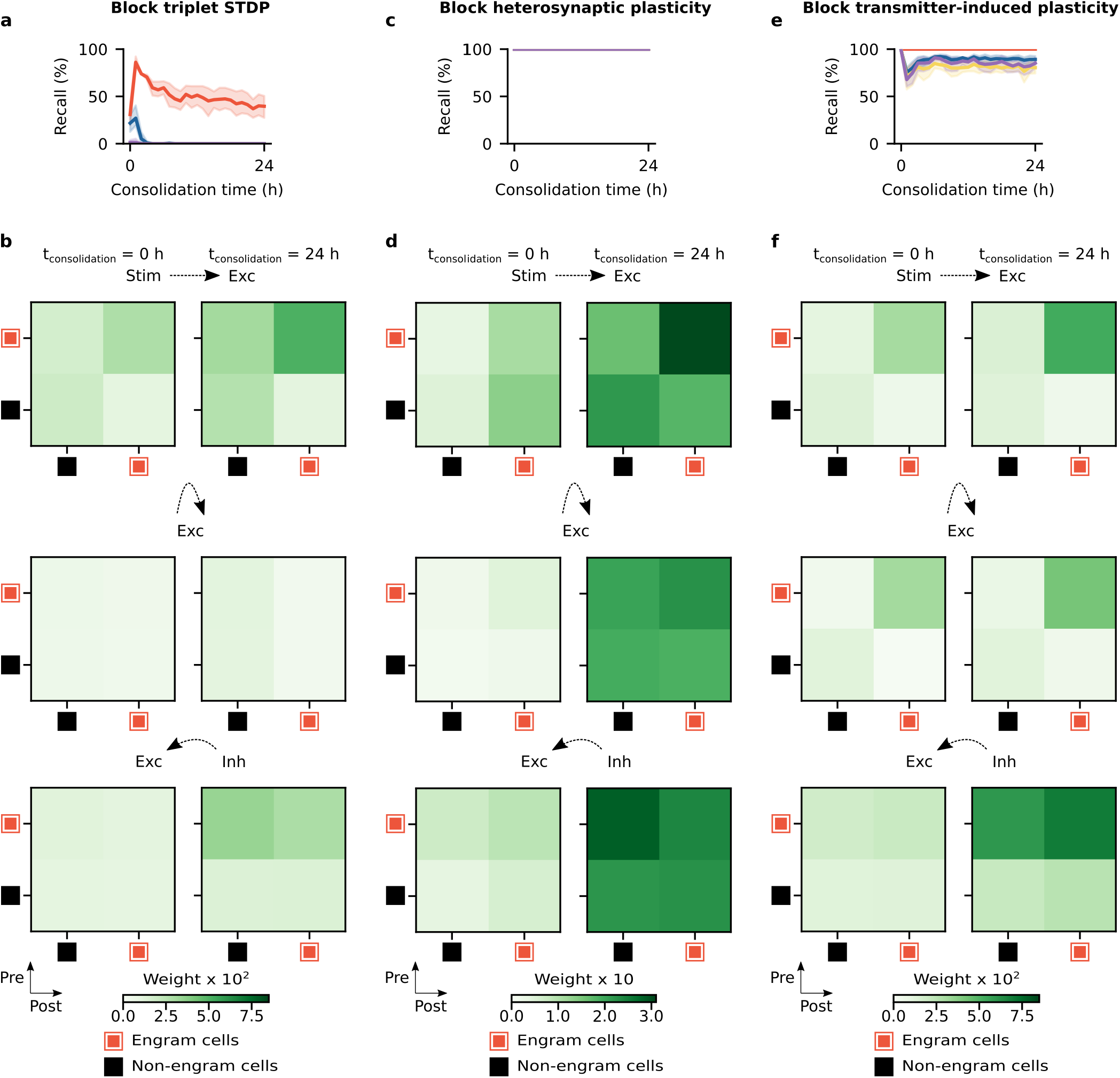
Hebbian and non-Hebbian forms of excitatory synaptic plasticity are critical for memory encoding and consolidation. **a-b**, Simulation protocol as in Fig. 1b but with blocked triplet STDP (i.e., *A* = *B_i_*(*t*) = 0, see Methods). **a**, Memory recall as a function of consolidation time. Means and 99% confidence intervals shown. *n* = 10. Color denotes stimulus as in Fig. 1c. **b**, Mean weight strength of plastic synapses clustered according to engram cell status (i.e., engram and non-engram cells). Top: feedforward excitatory synapses onto excitatory neurons. Middle: recurrent excitatory synapses onto excitatory neurons. Bottom: recurrent inhibitory synapses onto excitatory neurons. Left: at the end of the training phase. Right: after 24 hours of consolidation. Representative trial shown. **c-d**, Same as **a-b** but with blocked heterosynaptic plasticity (i.e., *β* = 0, see Methods). **e-f**, Same as **a-b** but with blocked transmitter-induced plasticity (i.e., *δ* = 0, see Methods).

**Extended Data Fig. 7.**
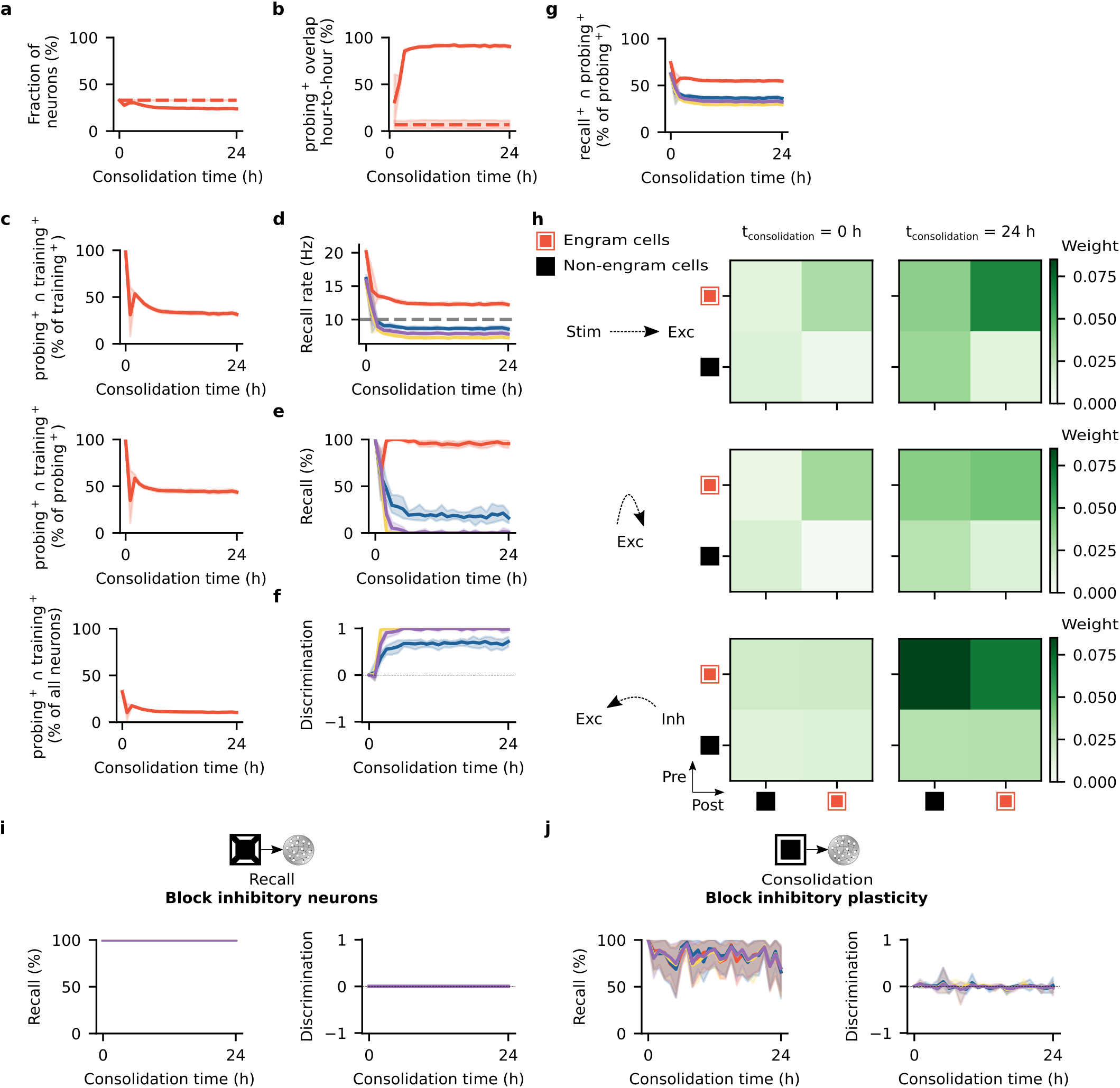
Alternative inhibitory synaptic plasticity can support consistent engram dynamics. **a-h**, Simulation protocol as in Fig. 1b but with the network subject to an alternative form of inhibitory synaptic plasticity (Equation 18, see Methods). **a-g**, Means and 99% confidence intervals shown. *n* = 10. **a**, Ensemble of engram cells as of fraction of all neurons (see Methods). Dashed line indicates engram cell ensemble at the end of training. **b**, Ensemble overlap between probing-activated engram cells at consolidation time = *t* and *t*-1 h as a fraction of engram cells at consolidation time = *t*-1 h. Dashed line indicates ensemble of neurons that remained part of the engram in all sampled time points (i.e., consolidation time = 0, 1, …, 24 h) as a fraction of engram cells at consolidation time = 0 h (i.e., training-activated engram cells). **c**, Ensemble overlap between engram cells activated during both probing and training as a fraction of training-activated engram cells (top), probing-activated engram cells (middle), and all neurons in the network (bottom). **d**, Firing rate of engram cells averaged across all cue presentations during recall as a function of consolidation time. Dashed line indicates threshold *ζ^thr^* = 10 Hz for engram cell activation. Color denotes stimulus as in Fig. 1c. **e**, Memory recall as a function of consolidation time. Color denotes stimulus as in Fig. 1c. **f**, Discrimination index between recall evoked by cues of the training stimulus and individual novel stimuli as a function of consolidation time (see Methods). Color denotes stimulus as in Fig. 1c. **g**, Fraction of probing-activated engram cells reactivated during recall as a function of consolidation time. Color denotes stimulus as in Fig. 1c. **h**, Mean weight strength of plastic synapses clustered according to engram cell status (i.e., engram and non-engram cells). Top: feedforward excitatory synapses onto excitatory neurons. Middle: recurrent excitatory synapses onto excitatory neurons. Bottom: recurrent inhibitory synapses onto excitatory neurons. Left: at the end of the training phase. Right: after 24 hours of consolidation. Representative trial shown. **i-j**, Means and 99% confidence intervals shown. *n* = 10. **i**, Simulation protocol as in Fig. 3a but with the network subject to an alternative form of inhibitory synaptic plasticity as in **a-h** (see Methods). Left: memory recall as a function of consolidation time. Right: discrimination index between recall evoked by cues of the training stimulus and individual novel stimuli as a function of consolidation time (see Methods). Color denotes stimulus as in Fig. 1c. **j**, Simulation protocol as in Fig. 4a but with the network subject to an alternative form of inhibitory synaptic plasticity as in **a-h** (see Methods). Left: memory recall as a function of consolidation time. Right: discrimination index between recall evoked by cues of the training stimulus and individual novel stimuli as a function of consolidation time (see Methods). Color denotes stimulus as in Fig. 1c. **a-j**, Network simulation parameters as in Fig. 1a-c except that 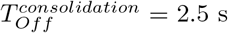 (see Methods and Supplementary Table 1).

**Extended Data Fig. 8.**
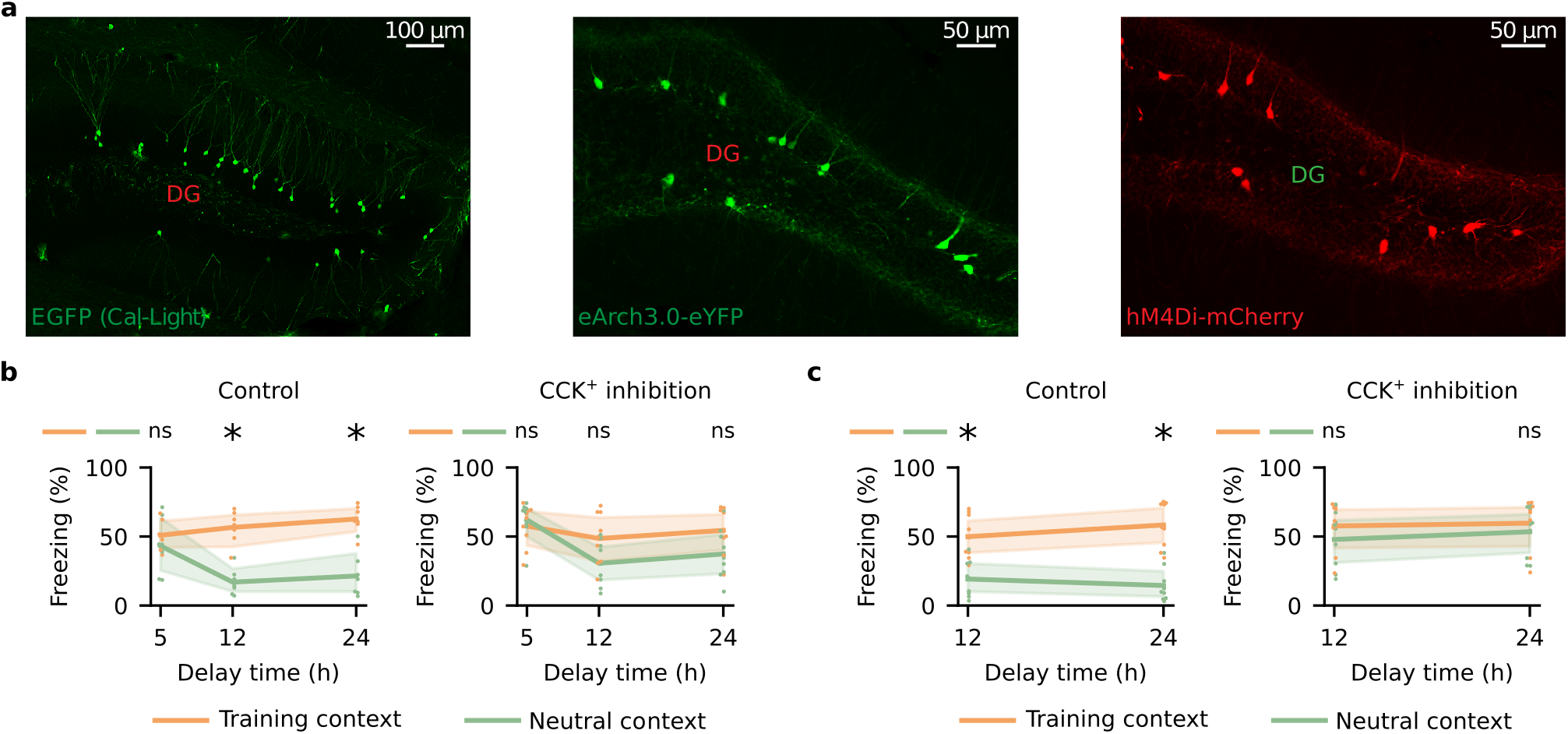
Representative images and freezing behavior in experiments. **a**, Representative images of experimental protocols. Left: Cal-Light labeling in the protocol in Fig. 2a. Middle: CCK^+^ neuronal eArch3.0-eYFP labeling in the protocol in Fig. 3e. Right: CCK^+^ neuronal hM4Di-mCherry labeling in the protocol in Fig. 4e. **b-c**, Means and 99% confidence intervals shown. **b**, Freezing levels during memory recall as a function of delay time in the protocol in Fig. 3e. Two-sided Wilcoxon signed-rank test between freezing in the training and neutral contexts. Control (*n* = 7 mice per group): for delay = 5 h, *W* = 11.0, *P* = 0.6875; for delay = 12 h, *W* = 0.0, *P* = 0.015625; and for delay = 24 h, *W* = 0.0, *P* = 0.015625. CCK^+^ inhibition (*n* = 9 mice per group): for delay = 5 h, *W* = 16.0, *P* = 0.496094; for delay = 12 h, *W* = 10.0, *P* = 0.164063; and for delay = 24 h, *W* = 8.0, *P* = 0.097656. **c**, Freezing levels during memory recall as a function of delay time in the protocol in Fig. 4e. Two-sided Wilcoxon signed-rank test between freezing in the training and neutral contexts. Control (*n* = 9 mice per group): for delay = 12 h, *W* = 0.0, *P* = 0.003906; and for delay = 24 h, *W* = 0.0, *P* = 0.003906. CCK^+^ inhibition (*n* = 9 mice per group): for delay = 12 h, *W* = 12.0, *P* = 0.25; and for delay = 24 h, *W* = 16.0, *P* = 0.496094. **b-c**, *: *P* < 0.05; ns: not significant.

**Extended Data Fig. 9.**
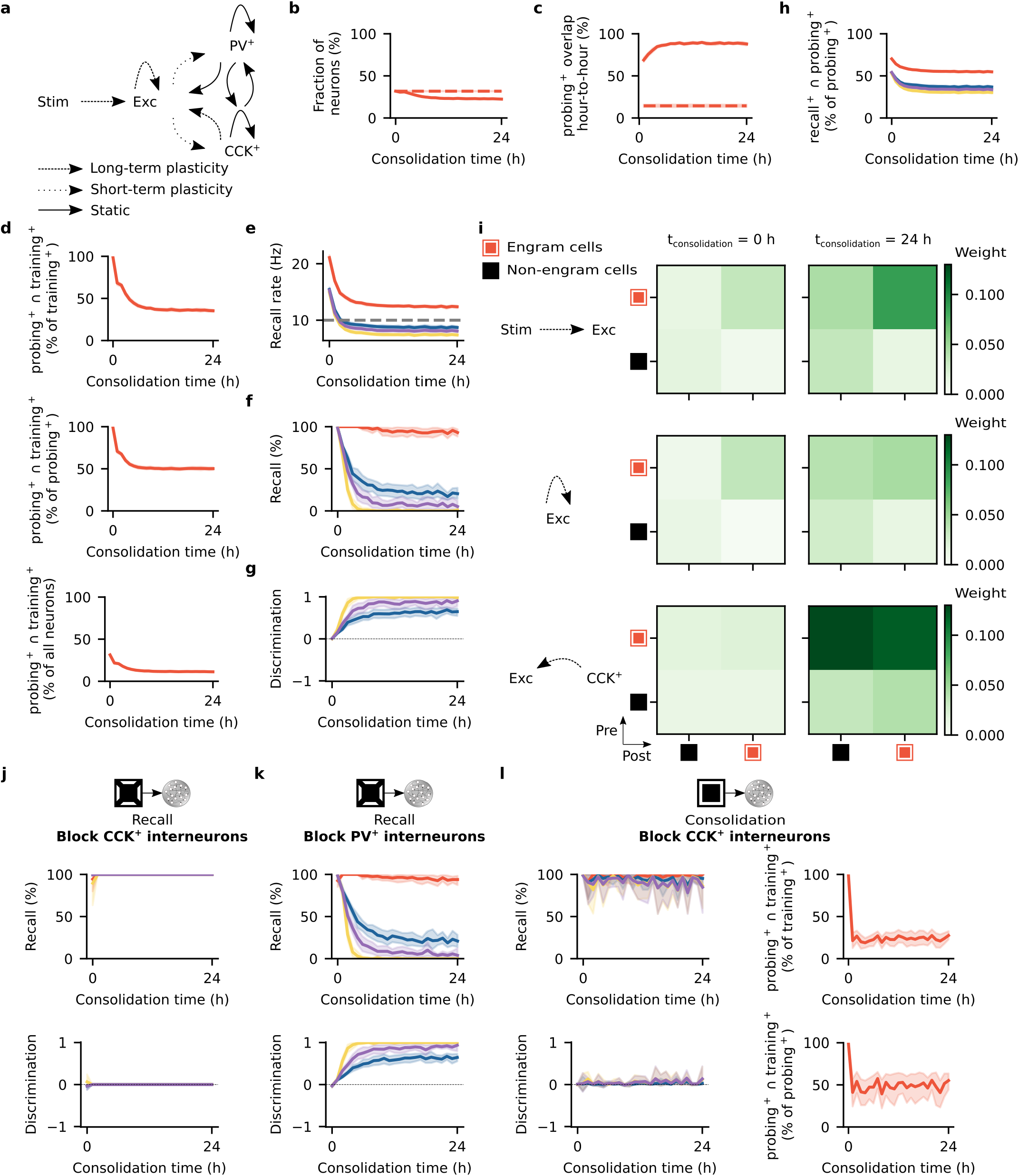
CCK^+^ interneurons support memory selectivity. **a**, Schematic of network model with CCK^+^ and PV^+^ interneurons. The network consists of a stimulus population (Stim) and a hippocampus network similar to Fig. 1a but with excitatory neurons (Exc) as well as CCK^+^ and PV^+^ interneurons. Plasticity of feedforward and recurrent synapses shown. Specifically, feedforward excitatory synapses and recurrent excitatory synapses onto excitatory neurons exhibit short- and long-term excitatory synaptic plasticity whereas recurrent excitatory synapses onto CCK^+^ or PV^+^ interneurons only display short-term plasticity. CCK^+^ synapses onto excitatory neurons are subject to inhibitory synaptic plasticity (Equation 15) while the remaining inhibitory synapses are static. For a detailed description of each form of plasticity, see Methods. **b-i**, Simulation protocol as in Fig. 1b with the network in **a**. **b-h**, Means and 99% confidence intervals shown. *n* = 10. **b**, Ensemble of engram cells as of fraction of all neurons (see Methods). Dashed line indicates engram cell ensemble at the end of training. **c**, Ensemble overlap between probing-activated engram cells at consolidation time = *t* and *t*-1 h as a fraction of engram cells at consolidation time = *t*-1 h. Dashed line indicates ensemble of neurons that remained part of the engram in all sampled time points (i.e., consolidation time = 0, 1, …, 24 h) as a fraction of engram cells at consolidation time = 0 h (i.e., training-activated engram cells). **d**, Ensemble overlap between engram cells activated during both probing and training as a fraction of training-activated engram cells (top), probing-activated engram cells (middle), and all neurons in the network (bottom). **e**, Firing rate of engram cells averaged across all cue presentations during recall as a function of consolidation time. Dashed line indicates threshold *ζ^thr^* = 10 Hz for engram cell activation. Color denotes stimulus as in Fig. 1c. **f**, Memory recall as a function of consolidation time. Color denotes stimulus as in Fig. 1c. **g**, Discrimination index between recall evoked by cues of the training stimulus and individual novel stimuli as a function of consolidation time (see Methods). Color denotes stimulus as in Fig. 1c. **h**, Fraction of probing-activated engram cells reactivated during recall as a function of consolidation time. Color denotes stimulus as in Fig. 1c. **i**, Mean weight strength of plastic synapses clustered according to engram cell status (i.e., engram and non-engram cells). Top: feedforward excitatory synapses onto excitatory neurons. Middle: recurrent excitatory synapses onto excitatory neurons. Bottom: recurrent CCK^+^ synapses onto excitatory neurons. Left: at the end of the training phase. Right: after 24 hours of consolidation. Representative trial shown. **j-l**, Means and 99% confidence intervals shown. *n* = 10. **j**, Simulation protocol as in Fig. 1b with the network in **a** but with CCK^+^ interneurons blocked during recall. Top: memory recall as a function of consolidation time. Bottom: discrimination index between recall evoked by cues of the training stimulus and individual novel stimuli as a function of consolidation time (see Methods). Color denotes stimulus as in Fig. 1c. **k**, Simulation protocol as in Fig. 1b with the network in **a** but with PV^+^ interneurons blocked during recall. Top: memory recall as a function of consolidation time. Bottom: discrimination index between recall evoked by cues of the training stimulus and individual novel stimuli as a function of consolidation time (see Methods). Color denotes stimulus as in Fig. 1c. **l**, Simulation protocol as in Fig. 1b with the network in **a** but with CCK^+^ interneurons blocked during consolidation. This also blocked the plasticity of CCK^+^ synapses onto excitatory neurons during consolidation (see Methods). The remaining CCK^+^ synapses continued to be static in the entire simulation protocol as in **a**. Left: memory recall as a function of consolidation time (top), and discrimination index between recall evoked by cues of the training stimulus and individual novel stimuli as a function of consolidation time (bottom) (see Methods). Color denotes stimulus as in Fig. 1c. Right: ensemble overlap between engram cells activated during both probing and training as a fraction of training-activated engram cells (top) and probing-activated engram cells (bottom). **a-l**, Network simulation parameters as in Fig. 1a-c except that 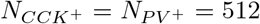 interneurons, 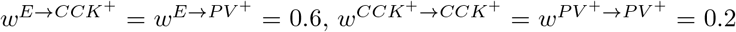,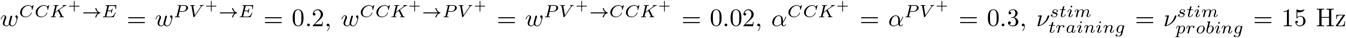,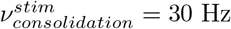, and 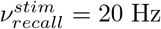 (see Methods and Supplementary Table 1).

## Supplementary Information

**Supplementary Table 1.** List of network simulation parameters. Value source indicated as below: (a) for values taken from Friedemann Zenke, Everton J Agnes, and Wulfram Gerstner. Diverse synaptic plasticity mechanisms orchestrated to form and retrieve memories in spiking neural networks. *Nature Communications*, 6(1):1–13, 2015. (b) for values chosen somewhat arbitrarily without targeted optimization. (c) for values optimized over several preliminary simulations.

| Neural Populations |  |  |
| --- | --- | --- |
| Parameter | Value<sup>(source)</sup> | Description |
| $N_{exc}$ | 4096<sup>(a)</sup> | Size of population of excitatory neurons in the hippocampus |
| $N_{inh}$ | 1024<sup>(a)</sup> | Size of population of inhibitory neurons in the hippocampus |
| $N_{stim}$ | 4096<sup>(a)</sup> | Size of population of stimulus neurons |
| Network Connectivity |  |  |
| Parameter | Value<sup>(source)</sup> | Description |
| $\epsilon_{rec}$ | 0.05<sup>(a)</sup> | Probability of connection of recurrent synapses |
| $R_{hpc}$ | 8<sup>(a)</sup> | Radius of receptive field of excitatory neurons |
| $w^{EE}$ | 0.1<sup>(a)</sup> | Initial weight of recurrent excitatory synapses onto excitatory neurons |
| $w^{EI}$ | 0.6<sup>(a)</sup> | Fixed weight of recurrent excitatory synapses onto inhibitory neurons |
| $w^{II}$ | 0.2<sup>(a)</sup> | Fixed weight of recurrent inhibitory synapses onto inhibitory neurons |
| $w^{IE}$ | 0.2<sup>(a)</sup> | Initial weight of recurrent inhibitory synapses onto excitatory neurons |
| $w_{stim}$ | 0.5<sup>(a)</sup> | Initial weight of feedforward excitatory synapses onto excitatory neurons |
| Neuron Model |  |  |
| Parameter | Value<sup>(source)</sup> | Description |
| $\tau^m$ | 20 ms<sup>(a)</sup> | Membrane time constant |
| $U^{rest}$ | -70 mV<sup>(a)</sup> | Membrane resting potential |
| $U^{exc}$ | 0 mV<sup>(a)</sup> | Excitatory reversal potential |
| $U^{inh}$ | -80 mV<sup>(a)</sup> | Inhibitory reversal potential |
| $\tau^{thr}$ | 5 ms<sup>(a)</sup> | Threshold time constant |
| $\vartheta^{rest}$ | -50 mV<sup>(a)</sup> | Threshold resting value |
| $\vartheta^{spike}$ | 100 mV<sup>(a)</sup> | Threshold value immediately after spike |
| Synapse Model |  |  |
| Parameter | Value<sup>(source)</sup> | Description |
| $\tau^{gaba}$ | 10 ms<sup>(a)</sup> | GABA decay time constant |
| $\tau^a$ | 100 ms<sup>(a)</sup> | Adaptation time constant |
| $\Delta^a$ | 0.1<sup>(a)</sup> | Adaptation strength |
| $\alpha^E$ | 0.2<sup>(a)</sup> | AMPA/NMDA ratio for excitatory neurons |
| $\alpha^I$ | 0.3<sup>(a)</sup> | AMPA/NMDA ratio for inhibitory neurons |
| $\tau^{ampa}$ | 5 ms<sup>(a)</sup> | AMPA decay time constant |
| $\tau^{nmda}$ | 100 ms<sup>(a)</sup> | NMDA decay time constant |
| Short-Term Plasticity Model |  |  |
| Parameter | Value<sup>(source)</sup> | Description |
| $\tau_{EE}^d$ | 150 ms<sup>(a)</sup> | Depression time constant for excitatory synapses onto excitatory neurons |
| $\tau_{EI}^d$ | 200 ms<sup>(a)</sup> | Depression time constant for excitatory synapses onto inhibitory neurons |
| $\tau^f$ | 600 ms<sup>(a)</sup> | Facilitation time constant for excitatory synapses |
| $U$ | 0.2<sup>(a)</sup> | Initial release probability for excitatory synapses |

| Long-Term Excitatory Synaptic Plasticity Model |  |  |
| --- | --- | --- |
| Parameter | Value<sup>(source)</sup> | Description |
| $\eta^{exc}$ | $1 \times 10^{-3(a)}$ | Learning rate of excitatory synapses |
| $\lambda^\beta$ | $50^{(a)}$ | $\beta/\eta^{exc}$ ratio |
| $\tau^{cons}$ | $20 \text{ min}^{(a)}$ | Synaptic consolidation time constant |
| $\lambda^\delta$ | $0.02^{(a)}$ | $\delta/\eta^{exc}$ ratio |
| $A$ | $1^{(a)}$ | LTP rate |
| $\tau^+$ | $20 \text{ ms}^{(a)}$ | Time constant of presynaptic trace for excitatory synaptic plasticity |
| $\tau^-$ | $20 \text{ ms}^{(a)}$ | Time constant of postsynaptic trace for excitatory synaptic plasticity |
| $\tau^{slow}$ | $100 \text{ ms}^{(a)}$ | Time constant of slow postsynaptic trace for excitatory synaptic plasticity |
| $P$ | $20^{(a)}$ | Potential strength |
| $w^P$ | $0.5^{(a)}$ | Upper fixed point of reference weight potential |
| $\tilde{w}$ | $0.0^{(a)}$ | Initial reference weight |
| $\tau^{hom}$ | $10 \text{ min}^{(a)}$ | Time constant of homeostatic regulation |
| $\tau^{ht}$ | $100 \text{ ms}^{(a)}$ | Time constant of postsynaptic trace for homeostatic regulation |
| $w_{exc}^{max}$ | $5.0^{(a)}$ | Maximum excitatory synaptic weight |
| $w_{exc}^{min}$ | $0.0^{(a)}$ | Minimum excitatory synaptic weight |
| Inhibitory Synaptic Plasticity Model |  |  |
| Parameter | Value<sup>(source)</sup> | Description |
| $\lambda^\eta$ | $50^{(a)}$ | $\eta^{exc}/\eta^{inh}$ ratio |
| $\gamma$ | $4 \text{ Hz}^{(a)}$ | Target activity level for excitatory neurons in the hippocampus |
| $\tau^{iSTDP}$ | $20 \text{ ms}^{(a)}$ | Time constant of pre- and postsynaptic traces for inhibitory synaptic plasticity |
| $\tau^H$ | $10 \text{ s}^{(a)}$ | Time constant of global secreted factor |
| $w_{inh}^{max}$ | $5.0^{(a)}$ | Maximum inhibitory synaptic weight |
| $w_{inh}^{min}$ | $0.0^{(a)}$ | Minimum inhibitory synaptic weight |
| Stimulus Model |  |  |
| Parameter | Value<sup>(source)</sup> | Description |
| $\nu^{bg}$ | $5 \text{ Hz}^{(a)}$ | Background firing rate of stimulus population |
| $\nu^{stim}$ | $15 \text{ Hz}^{(c)}$ | Firing rate of neurons matching given stimulus when that stimulus is activated |
| $T_{On}^{training}$ | $1 \text{ s}^{(c)}$ | Mean stimulus-on period in the training phase |
| $T_{Off}^{training}$ | $2 \text{ s}^{(c)}$ | Mean stimulus-off period in the training phase |
| $T_{On}^{probing}$ | $1 \text{ s}^{(c)}$ | Mean stimulus-on period in the probing phase |
| $T_{Off}^{probing}$ | $2 \text{ s}^{(c)}$ | Mean stimulus-off period in the probing phase |
| $T_{On}^{consolidation}$ | $1 \text{ s}^{(c)}$ | Mean stimulus-on period in the consolidation phase |
| $T_{Off}^{consolidation}$ | $3 \text{ s}^{(c)}$ | Mean stimulus-off period in the consolidation phase |
| $T_{On}^{recall}$ | $1 \text{ s}^{(c)}$ | Mean stimulus-on period in the recall phase |
| $T_{Off}^{recall}$ | $2 \text{ s}^{(c)}$ | Mean stimulus-off period in the recall phase |
| Network Simulation |  |  |
| Parameter | Value<sup>(source)</sup> | Description |
| $T_{burn}$ | $120 \text{ s}^{(b)}$ | Duration of burn-in period prior to training |
| $T_{training}$ | $5 \text{ min}^{(c)}$ | Duration of training phase |
| $T_{consolidation}$ | $24 \text{ h}^{(b)}$ | Duration of consolidation phase |
| $T_{probing}$ | $60 \text{ s}^{(c)}$ | Duration of probing phase |
| $T_{recall}$ | $90 \text{ s}^{(b)}$ | Duration of recall phase |
| $\Delta$ | $0.1 \text{ ms}^{(a)}$ | Time step for updating neuronal state variables except for reference weights $\tilde{w}$ |
| $\Delta_{long}$ | $1.2 \text{ s}^{(a)}$ | Time step for updating reference weights $\tilde{w}$ |
| $N_{ranks}$ | $16^{(c)}$ | Number of MPI ranks in Auryn simulations |

